# Cardiac fibrosis inhibitor CTPR390 prevents structural and morphological changes in collagen and fibroblasts of engineered human connective tissue

**DOI:** 10.1101/2025.01.15.633162

**Authors:** David Maestro, Ana Palanca, Helena Soto, I Llarena, Alisa Nicole DeGrave, Gabriela Guedes, Guilherme Henrique de Oliveira, André Luiz Coelho Conceição, Verónica Mieites, Jose M. Icardo, Carlos Sanchez-Cano, Olga M. Conde, Susanne Lutz, Aitziber L. Cortajarena, Ana V. Villar

## Abstract

Cardiac fibrosis is a key characteristic of heart failure, with no effective treatment available. Using three-dimensional human models and cutting-edge biotechnology to evaluate new therapies offers a significant advancement. CTPR390, an experimental anti-fibrotic inhibitor targeting Hsp90, has shown success in animal models, but remains unexplored in human cardiac models. This study evaluated a cardiac three-dimensional engineered connective tissue (ECT) model treated with CTPR390, focusing on changes in the extracellular matrix and fibroblasts. Results showed that CTPR390 prevented architectural changes in TGFβ1-activated ECT, preserving tissue perimeter, collagen fibers alignment while reducing percentage of structured areas and degree of collagen structuration. Additionally, the treatment reduced cell area of elongated fibroblasts under tension, without changes in the internal rounded cells devoid of tension. Fibroblast recruitment to tension areas was diminished, showing biomechanical behavior similar to control ECT. This treatment also lowered the gene and protein expression of key pro-fibrotic markers. For the first time, advanced biotechnology was employed to detect the detailed structure of tissue fibrosis reduction after administering CTPR390, representing a significant advancement toward clinical application for cardiac fibrosis treatment.

**Graphical Abstract:** 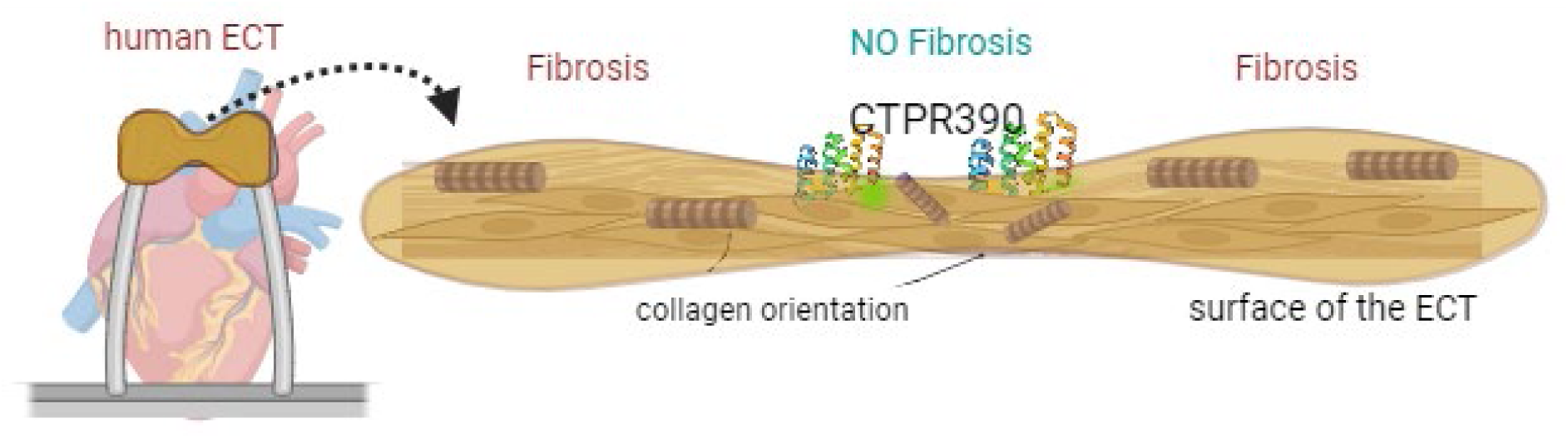

## Introduction

Cardiovascular disease remains the leading cause of death in industrialized countries, significantly impacting morbidity and mortality rates. Cardiac fibrosis, a common secondary condition in various cardiovascular diseases, is characterized by excessive extracellular matrix (ECM) protein deposition, which, while initially protecting the heart after myocardial damage, ultimately leads to functional loss. Among the three main types of cardiac fibrosis: replacement, perivascular and interstitial fibrosis [1], interstitial fibrosis can occur in response to various cardiovascular conditions such as aortic valve dysfunction or hypertension. Although remodeling initially helps maintain cardiac function, unchecked fibrosis can lead to excessive ECM buildup, increasing stiffness and contributing to heart failure. Therefore, the extent of fibrosis is considered a predictor of adverse outcomes in heart disease patients [2]. Despite extensive research, effective therapies for cardiac fibrosis are lacking [3]. A major challenge is the absence of human cardiac models that accurately replicate human heart disease features [4]. Animal models while offering valuable insights into remodeling patterns [5–7], have limitations compared to humans. Engineered heart models created from human stem cell, such as engineered human myocardium (EHM) [8], and human engineered heart tissue (EHT) [9, 10] and simpler models like engineered connective tissue (ECT) provide valuable insights [11, 12]. ECT models, incorporating human cardiac fibroblasts in collagen hydrogels, effectively mimic the 3D architecture of adult cardiac fibroblasts, making them ideal for studying fibrotic diseases [13]. Understanding ultrastructural changes in the fibrotic ECM, such as fiber alignment [14], and collagen scar [15], is crucial for evaluating advanced treatments. This study employs ECT to explore mechanical, biochemical, and ultrastructural characteristics of fibroblasts and ECM fibers exposed to an experimental anti-fibrotic nanodrug. ECT fibrosis is induced by Transforming Growth Factor β (TGFβ) [16], a cytokine secreted by fibroblasts [17]. TGFβ initiates the pro-fibrotic process [18], and activates its receptor complex on the cell surface. Disrupting TGFβ signaling in cardiac fibroblasts has been linked to fibrosis reduction [19, 20]. Heat shock protein 90 (Hsp90), an ATP-dependent molecular chaperone, aids protein folding. Inhibiting Hsp90, particularly its C-terminal end, can reduce TGFβ signaling activity [21]. These approaches include the reduction of TGFβ signaling activity by disrupting the TGFβ receptor complex involving Hsp90 [19]. One such C-terminal end Hsp90 inhibitor tested as an anti-fibrotic molecule is the engineered consensus tetratricopeptide repeat inhibitor of Hsp90 (CTPR390) [19, 22, 23]. The protein region of CTPR390 (ProtCTPR390) consists of three consensus tetratricopeptide repeat (CTPR) units repeated in tandem, which were engineered drawing inspiration from natural binding domains like tetratricopeptide repeats (TPR) found in proteins such as Hsp70/Hsp90 organizing protein (HOP) [24]. Through genetic manipulation, ProtCTPR390 was combined with other functional motif, i.e. a gold nanocluster (AuNC) coordinating domain [25] designed to display cysteines at strategic locations, creating multifunctional chimeric proteins [26, 27]. These proteins were additionally conjugated with Alexa-488, resulting in stable, trackable constructions under biological conditions through nanocluster or fluorophore-specific imaging while harboring the active module capable of binding to Hsp90 [28–30]. The binding occurs at the C-terminal end of Hsp90, thus inhibiting the interaction of Hsp90 with the TGFβ receptor complex, which disrupts the proper functioning of the TGFβ-dependent fibrosis signaling cascade [19, 22].

The TGFβ-activated ECT from cardiac patient fibroblasts offers a valuable model to explore biomechanical and biochemical precise responses to advanced anti-fibrotic drugs like CTPR390. Additionally, it presents a unique opportunity to reveal, for the first time in a human 3D model and using highly advanced technology, various ultrastructural features of human cardiac collagen and fibroblast behavior following an anti-fibrotic action.

## Results

### CTPR390 restores physical and biomechanical properties of fibrotic ECTs

We generated ECT composed of primary adult cardiac fibroblasts and bovine collagen I within molds equipped with flexible poles (Figure 1a). Polarization-Sensitive Optical Coherence Tomography (PS-OCT) provided, for the first time, detailed images of the ECT structure showing both top-down and side views (Figure 1a). Three groups of ECTs were cultivated for 13 days: Control, TGFβ1 (to induce fibrosis) and CTPR390 to prevent fibrosis. CTPR390 was engineered with an Hsp90 binding module and Alexa 488 fluorophore for visualization, and its purity, stability (Supplementary Figure 1a) and molecular weight (Supplementary Figure 1a-b) were confirmed. The folding, stability signatures of CTPR390-AuNC (Supplementary Figure 1c) and the emission spectra of CTPR90-AuNC-488 named CTPR390 (Supplementary Figure 1c-f) were also verified. To assess physical characteristics, we generated dispersion maps comparing conditions. Control/CTPR390 ECTs had a higher overlap in compactness (64.5%) compared to Control/TGFβ1 (56.9%) and CTPR390/TGFβ1 (40.7%) (Figure 1b). Fiber orientation analysis revealed that CTPR390 prevented fibrosis, producing fiber alignment similar to the control with fewer fibers at angles above 120°, while TGFβ1 led to highly structured fibers more aligned at angles below 60° (Figure 1c-d) (Supplementary Figure 2a). We also measured tissue organization along depth (z), with TGFβ1 showing a notable decline in fiber alignment, which was corrected by CTPR390, bringing it closer to control levels (Figure 1e). Biomechanical analysis over 13 days showed increased contraction in TGFβ1-treated ECTs from day 7, while CTPR390 prevented these changes, maintaining control group behavior (Figure 1f-g). The Young’s Modulus, reflecting ECT stiffness, was determined from the stress-strain curves and those measurements unveiled a significant increase in stiffness of TGFβ1 group compared to Control group (***p < 0.0005), which was prevented by co-application of CTPR390 (****p < 0.0001) (Figure 1h). In a similar fashion, the administration of CTPR390 resulted in an inhibition of the TGFβ1-dependent decline of the capacity of the ECT to absorb elastic strain energy before yielding (resilience) (Figure 1i). These findings suggest that the TGFβ1-activated ECT treated with CTPR390 exhibited elastic properties and a recovery capacity from deformation similar to those observed under control condition (Figure 1i).

**Figure 1:**
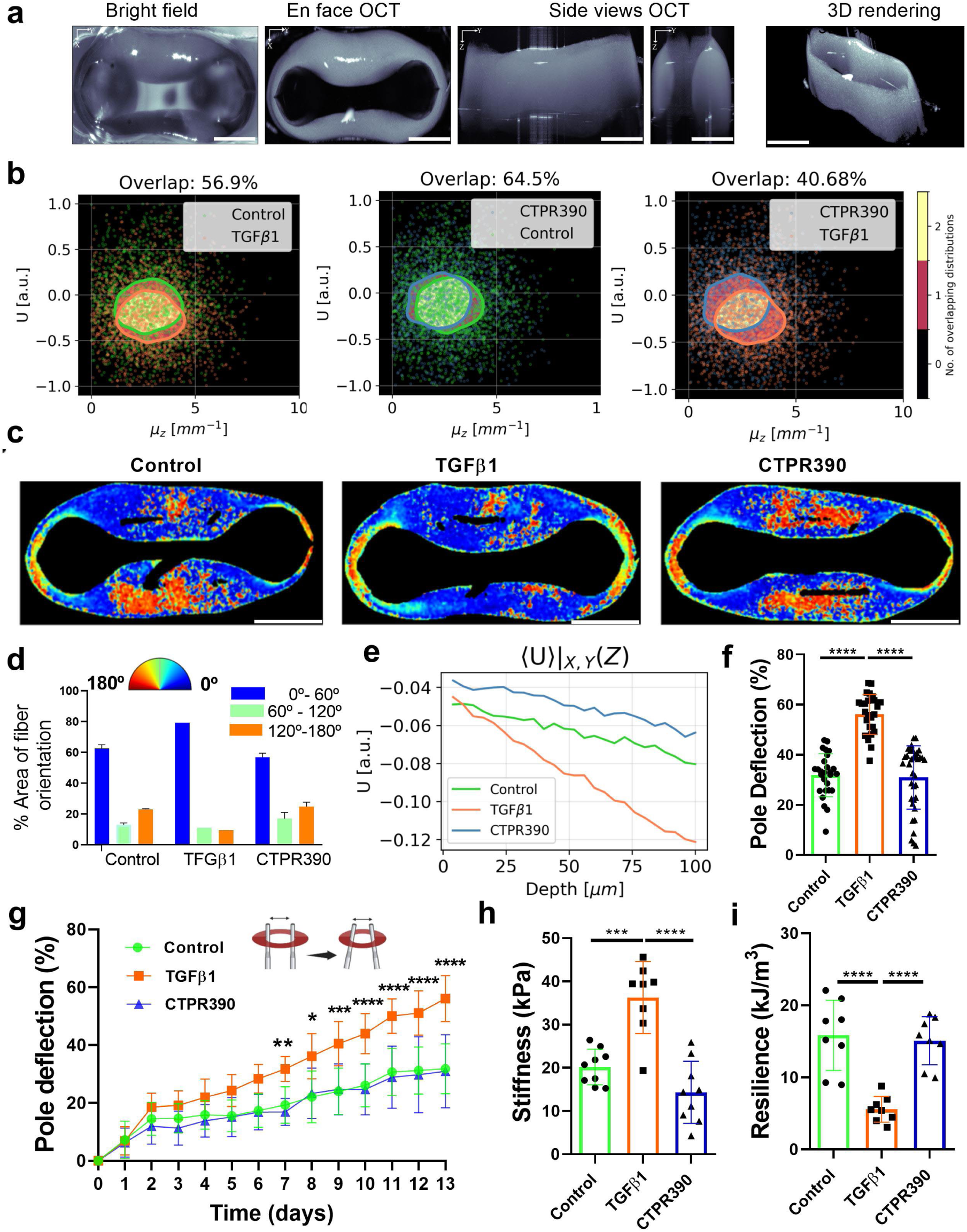
Administering CTPR390 to TGFβ1-activated ECT resulted in a reversal of its physical and biomechanical characteristics to those observed in control ECT. **a)** Representative images (bright field image, en face view, side views, and 3D rendering) of an ECT using PS-OCT technique that showed the morphology of the engineered tissue around the two flexible poles. The white scale bar indicated 1.3 mm. **b)** Three panels showing the correlative behavior observed within a sigma area between the Stokes U and the attenuation coefficient (μm^-1^), considering the quantified overlap in distributions in Control, TGFβ1, and CTPR390 ECT samples. n = 3 biological samples (ECT) measuring 10⁸ pixels in triplicate distributed in layers of 3 3.6 um/px thick, assuming a refractive index of n=1.38, typical of biological samples, evaluated up to the first 100 micrometers. White scale bar indicated 1 mm. **c)** Representative ECT images of the angle of polarization (AoP) orientation aligned with the fibers within the ECT using the same color-coding as in panel. n = 3 biological samples (ECT) measuring 10⁸ pixels in triplicate distributed in layers of 3 3.6 um/px thick, assuming a refractive index of n=1.38, typical of biological samples, evaluated up to the first 100 micrometers in Control, TGFβ1 and CTPR390 groups. The white scale bar indicated 1.3 mm. **d)** Bar graph illustrating the quantification of areas with the same percentage of AoP aligned with ECT fibers, using the PS-OCT technique. AoP color code: blue (0° – 60°), green (60° – 120°), and orange (120° – 180°). The process from data acquisition segmenting areas with the same AoP and assigning the color code for the AoP is detailed in Supplementary Figure 2a. n = 3 biological samples (ECT) measuring 10⁸ pixels in triplicate distributed in layers of 3 3.6 um/px thick, assuming a refractive index of n=1.38, typical of biological samples, evaluated up to the first 100 micrometers **e)** Evolution along depth (z) of the xy-averaged Stokes U parameter (a.u.) among the Control, TGFβ1, and CTPR390 groups within the initial 100 μm from the surface of the ECT. Control Y = (−554.8e-04 ± 8.0e-04) + (−129.8e-05 ± 5.7e-05) X. TGFβ1 Y = (−525.7e-04 ± 8.1e-04) + (−362.1e-05 ± 5.8e-05) X. CTPR390 Y = (−431.9e-04 ± 8.0e-04) + (−138.3e-05 ± 5.7e-05) X. n = 3 biological samples (ECT) measuring 10⁸ pixels in triplicate distributed in layers of 3 3.6 um/px thick, assuming a refractive index of n=1.38, typical of biological samples, evaluated up to the first 100 micrometers. **f)** Detailed analysis of contraction at day 13 presented in a bar graph comparing Control and CTPR390, to TGFβ1 group. Values presented as means ± SEM for n = 12 ECT per group with triplicates of each biological sample (**** p < 0.0001) were assessed through 2-way ANOVA with Tukey’s multiple comparison test. **g)** ECT contraction was evaluated over a 13-day period based on the percentage (%) of pole deflection in three groups (Control, TGFβ1, and CTPR390), n = 12 ECT per group with triplicates of each biological sample. **h)** Bar graph illustrating ECT stiffness (Young’s modulus) with significant differences observed in TGFβ1 compared to Control or CTPR390 groups n = 8 - 9 ECT per group with triplicates of each biological sample. **i)** Bar graph depicting ECT resilience with significant differences noted in TGFβ1 compared to Control or CTPR390 groups. n=7-9 ECTs with triplicates of each biological sample. P values (*p < 0.05, **p < 0.005, ***p < 0.0005, ****p < 0.0001) were determined using one-way ANOVA with Tukey’s multiple comparison test.

### CTPR390 reduces pro-fibrotic gene and protein expression in ECTs

Previous data from our group detailed the anti-fibrotic mechanism of action of the CTPR390, which inhibits Hsp90 by binding to its C-terminal end. This causes a conformational change in the Hsp90 dimer and displaces it from the TGFβ receptor complex, altering TGFβ signaling without toxicity when CTPR390 enters fibroblasts [19, 31]. This activity reduced the expression of collagen and other pro-fibrotic markers [19, 22, 23]. Building on these findings, we demonstrated anti-fibrotic effects of CTPR390 in TGFβ1-activated ECT for the first time. After administering CTPR390 to ECTs with TGFβ1-activated fibroblasts, fluorescence from Alexa-488 confirmed the presence of CTPR390 throughout the tissue (Figure 2a-b). Further examination revealed CTPR390 inside human fibroblasts (Figure 2c). An enzymatic digestion process was then employed to extract fibroblasts from the ECT, followed by cultivation to re-confirm the presence of CTPR390 and assess cell survival in culture after 13 days within the ECT (Figure 2d-e). Notably, a significant percentage of the total fibroblasts within the ECT demonstrated internalization of CTPR390 (83.3 % ± 7.0 %) (Figure 2d). This incorporation did not induce cell toxicity, as evidenced by the comparable viability of cells treated and non-treated with CTPR390 (Supplementary Figure 2b).

**Figure 2:**
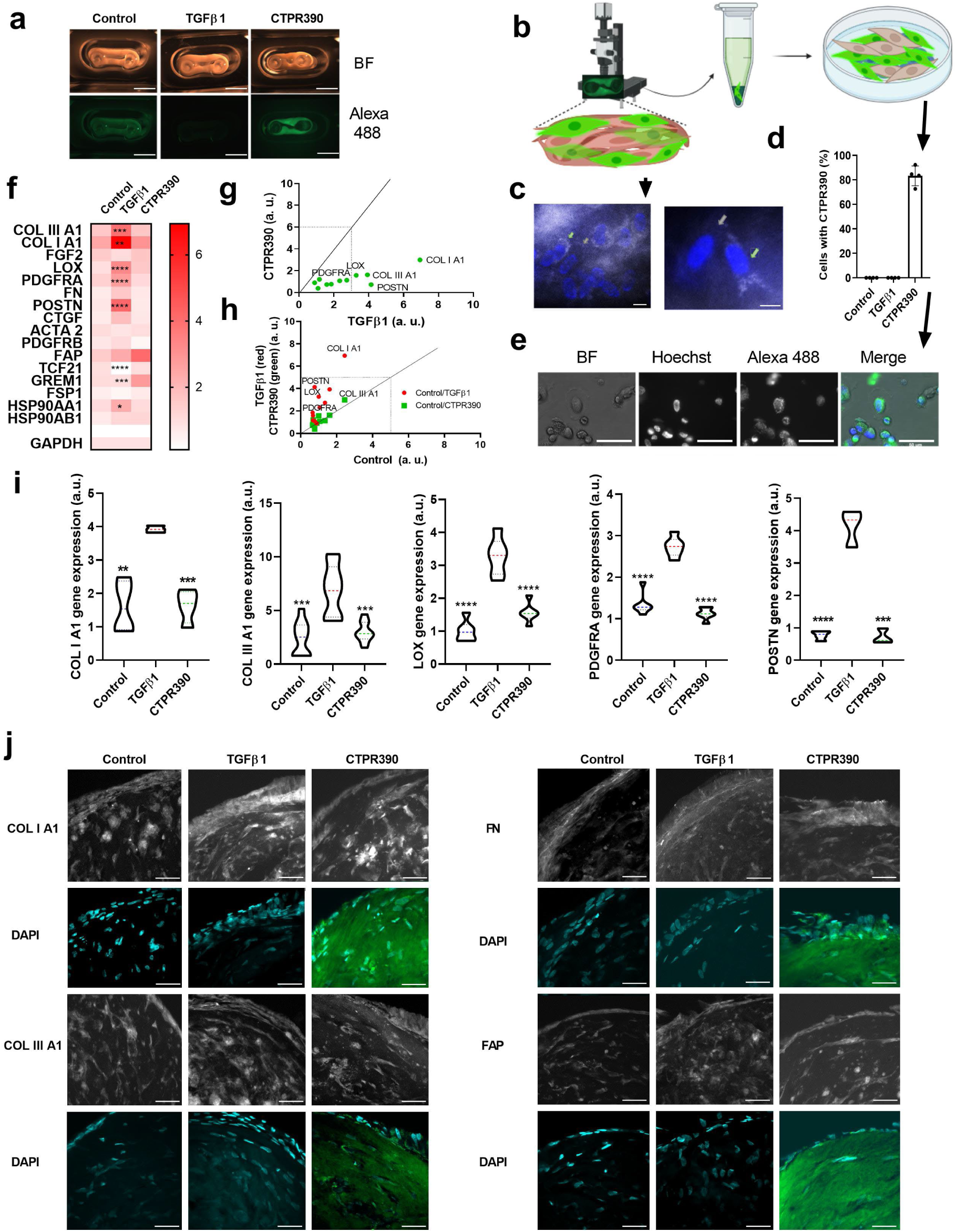
Restoration of normal ECT gene and protein expression following the introduction of CTPR390 to TGFβ1-activated ECT. **a**) Bright-field and epifluorescence imaging of control representative images of ECT, TGFβ1-activated ECT, and TGFβ1-activated ECT subjected to CTPR390 treatment (Control, TGFβ and CTPR390). Both white scale bars in the two panels indicated 5 mm. **b**) Diagram illustrating the detection of CTPR390-green fluorescence (resulting from Alexa-488 fluorophore conjugated to CTPR390) incorporated into ECT. It included a close-up view of the fluorescent fibroblasts within the ECT and the subsequent extraction of fibroblasts from the ECT, transferred to a tube, and later seeded onto a culture plate, revealing both fluorescent and non-fluorescent fibroblasts. Diagram generated with Biorender. **c**) CTPR390 fluorescence captured directly from the surface of the TGFβ1 ECT following a 13-day treatment with CTPR390. The white scale bar indicated 10 μm. **d**) Bar graphs presenting the percentage and total cells retaining CTPR390 over a 13-day period in CTPR390 ECT, n = 4 ECT per group, with three technical replicates. **e**) Visualization of bright-field, nuclei (Hoechst) and CTPR390 (Alexa-488), along with the merged representation of all three images, pertaining to purified fibroblasts derived from CTPR390 group and cultured in 2D plates. The white scale bar indicated 50 μm. **f**) Heatmap illustrating the prevalent pro-fibrotic genes, with darker red representing higher expression (2x over the median) and light reds indicating lower expression (0.5 lower the median) in Control, TGFβ1, and CTPR390 ECT groups; n = 6-10 ECT per group with 3 technical replicates of each ECT sample. **g-h**) XY plots illustrating the comparison of expression for all pro-fibrotic genes tested between CTPR390 and TGFβ1 (**g),** as well as between TGFβ1 and Control groups (**h**). **i**) Bar graphs illustrating significant gene expression differences of the most differentially expressed pro-fibrotic genes tested in Control, TGFβ1 and CTPR390 groups; n = 6-10 ECT per group with 3 technical replicates of each ECT sample. The p values (**p < 0.005, ***p<0.0005, ****p < 0.0001) were determined using a one-way ANOVA with Tukey’s multiple comparison test. **j**) Representative confocal images depicting the localization of COL I A1, COL III A1, FN, and FAP proteins in cross-sections of ECTs. A higher density of superficial cells (indicated by DAPI-stained nuclei), and COL IA1, was observed in TGFβ1-activated ECTs compared to the Control and CTPR390-treated groups. CTPR390 was visualized using Alexa 488 fluorescence. The white scar bars indicated 50 μm.

Gene expression analysis of myofibroblast and fibroblast markers in TGFβ1-activated ECT [16], showed significant differences, illustrated in a heatmap (Figure 2f). Two xy plots were used to identify the most differentially expressed genes by comparing the Control ECT group with the TGFβ1 group and the TGFβ1 group with the CTPR390 group (Figure 2g-h). We emphasized the potent anti-fibrotic effect of CTPR390 by demonstrating a significant reduction in the expression of critical pro-fibrotic genes, clearly showing differences in the gene expression of COL I A1, COL III A1, PDGFRA, LOX, and POSTN, (Figure 2i). At the protein level, we observed a marked reduction in COL I A1 expression and a slight decrease in FAP in CTPR390-treated fibroblasts compared to TGFβ1-activated fibroblasts. In all cases, CTPR390 treatment reduced the number of superficial cells, along with key pro-fibrotic protein expression (COL IA1 and FAP) in these areas. No changes were detected in FN expression or COL III A1 (Figure 2j). Interestingly, the same human cardiac fibroblasts cultured in 2D showed no differences in COL I A1 and FAP protein expression between Control and TGFβ1-activated fibroblasts, except for COL III A1 (Supplementary Figure 2c).

### CTPR390 avoids the pro-fibrotic cell profile in TGFβ1-treated ECT

In all groups (Control, TGFβ1, and CTPR390), hematoxylin-eosin staining of cross-sections extracted from the medial region of the long branch of the ECT revealed elongated superficial fibroblasts compared to inner cells (Figure 3a). Surface cells were aligned in the pole-to-pole direction, while interior cells were arranged randomly within the ECT (Figure 3a). CTPR390 treatment significantly reduced the number of fibroblast layers on the surface of TGFβ1-activated ECT, from 4.5 ± 0.5 layers to 2.5 ± 0.5 layers (**p < 0.005), bringing it closer to control levels (Figure 3b). Scanning electron microscopy showed higher surface activity in TGFβ1-activated ECT, with more filopodia and blebs, indicating intense cell-matrix interactions and membrane stress. In contrast, control and CTPR390-treated ECTs exhibited less cellular activity, with reduced filopodia presence (Figure 3c). The observed bleb formation in TGFβ1-activated fibroblasts of fibrotic ECT is typically seen when the cell membrane detaches from the underlying actin cytoskeleton [32]. This can occur due to increased internal pressure and mechanical stress, causing the membrane to balloon outward [33]. In contrast, control and CTPR390-treated ECT showed less cellular activity, although CTPR390 treatment still maintained some active surface areas by showing reduced areas with filopodia (Figure 3c). Cell count analysis showed a reduction in cell area (Figure 3d) and cell perimeter (Figure 3e) in CTPR390-treated ECTs compared to TGFβ1-activated ones (Figure 3d-e). Fibroblasts from the ECT, as happens in the heart [34], appeared to be a key component for sustaining the tension of the tissue. To corroborate this, we used collagen without fibroblasts to form an ECT (Supplementary Figure 3a), which resulted in the formation of a diffused, not solidified ECT unattached to the poles. However, the presence of fibroblasts resulted in a tense ECT that was perfectly attached to the two poles and presented regular short and long arms (Supplementary Figure 3b). Moreover, when collagen was deposited and solidified in culture media, it formed a dense, uniform layer from any angle of observation, indicating the absence of varying angles of polarization (AoP) (Supplementary Figure 3c-d). Lysotracker labeling confirmed higher cell density on the surface of TGFβ1-activated ECT compared to Control (****p < 0.0001) and CTPR390-treated ECT (*p < 0.05) (Figure 3f-g). The heightened activation state of fibroblasts in TGFβ1-activated ECT was supported by increased actin cytoskeleton abundance [35] stained with phalloidin (Figure 3h). In contrast, CTPR390-treated ECT showed normal actin expression and reduced surface cell accumulation, as confirmed by nuclear DAPI staining (Figure 3h).

**Figure 3:**
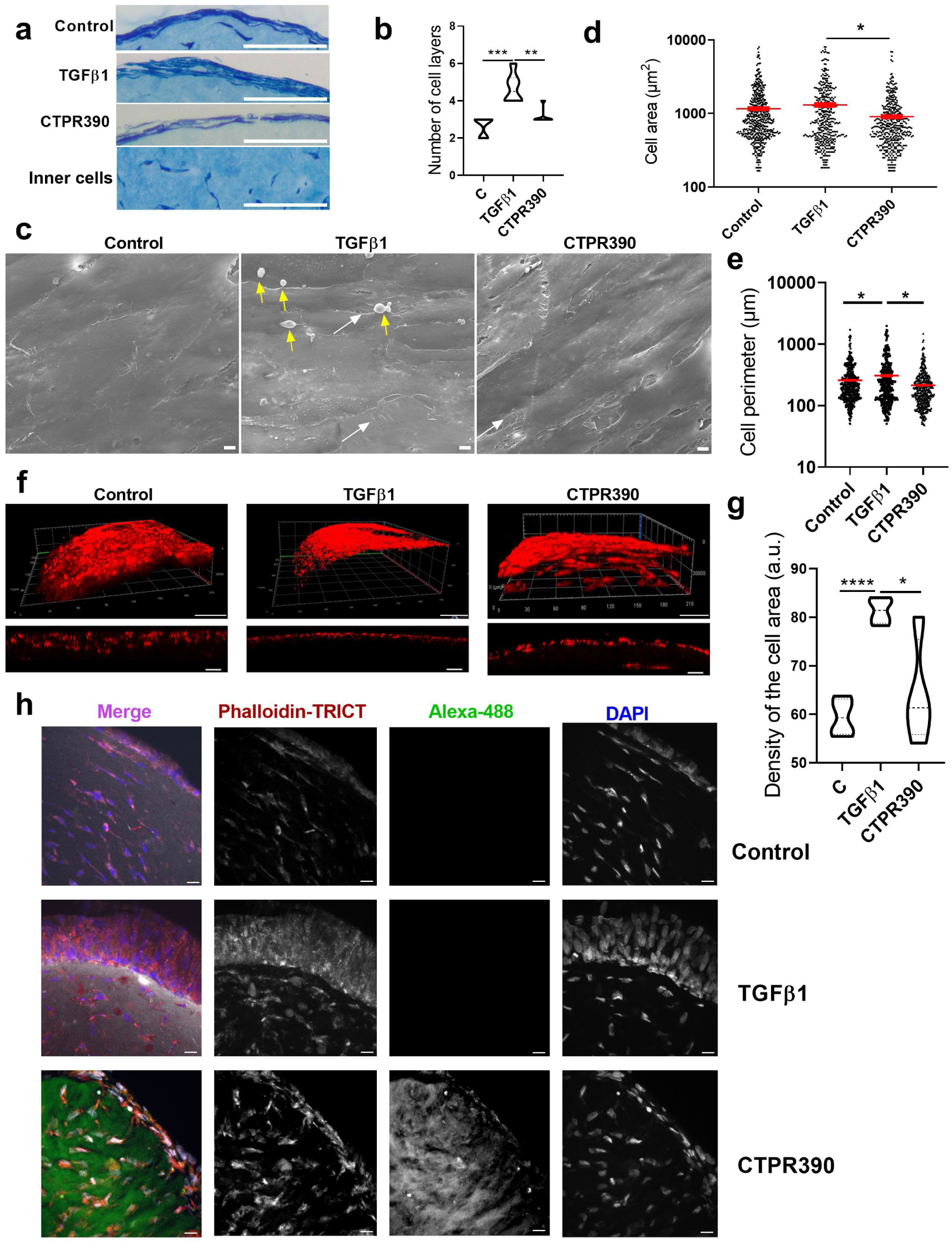
Attenuation of TGFβ1-activated ECT pro-fibrotic cell features after CTPR390 treatment. **a)** Representative images of ECT surface with fibroblasts stained with hematoxylin-eosin from cross-sections of Control, TGFβ1 and CTPR390 ECTs and representative image of inner fibroblast from an ECT; the white scale bars indicated 50 μm. **b**) Violin plots depicting significant differences in the number of cell surface layers of TGFβ1 ECT compared to Control and CTPR390 ECTs, n = 6 independent samples measured in triplicate. **c**) Three panels featuring a depiction with a front view of fibroblasts on the surface of Control, TGFβ1 and CTPR390 ECTs by scan microscopy of 1 ECT per group. Yellow arrows indicate surface blebs, and white arrows indicate filopodia in TGFβ1and CTPR390 ECTs; the white scale bars indicated 2 μm. **d**) Cell area count and, **e**) Cell perimeter count in the three groups of ECT (Control, TGFβ1 and CTPR390) performed in n = 12 ECT per group, n of cells = 400-510 cells per group. **f**) Confocal images showcasing the 3D reconstruction of cellular surfaces within 100 μm depth utilizing lysotracker (in red) as a cell dye for comparing cell surface density in Control, TGFβ1, and CTPR390 ECTs. The white scale bars in upper panels indicated 30 μm and the white scale bars in lower panels indicated 40 μm. **g**) Violin graph presenting the analysis of significant differences in the density of cell area in Control, TGFβ1, and CTPR390 ECTs; n = 4 independent ECT samples measured in triplicate. The p values (*p < 0.05, **p < 0.005, ***p < 0.0005, ****p < 0.0001) of all these graphs were determined using a one-way ANOVA with Tukey’s multiple comparison test. **h**) Representative confocal images of ECT cross-sections visualizing human actin-F (phalloidin-TRICT staining), CTPR390 fluorescence (Alexa 488), and nuclei (DAPI staining) in the surface and inside areas of control, TGFβ1 and CTPR390 ECT. The white scale bars indicated 10 μm.

### CTPR390 preserves fibroblast distribution and structure in ECT

At day 0, 750,000 human primary cardiac fibroblasts are embedded in the bovine collagen to generate the 3D engineered tissue; once the ECT compacts on day 1, the model featured a media of 680,000 homogeneously distributed fibroblasts throughout the 3D tissue (Figure 4a-b). After 13 days of culture, the cell organization changed and exhibited a particular segregation with differential cell separation between the ECT surface and the ECT interior (Figure 4c), with a total cell number, presenting variations depending on the group: Control 247,000 ± 35,637; TGFβ1 234,000 ± 34,351; or CTPR390 232,500 ± 23,629 (Figure 4d). A reduction in the initial cell number due to failed cell attachment regularly occurred during the maturation process of the ECT.

**Figure 4:**
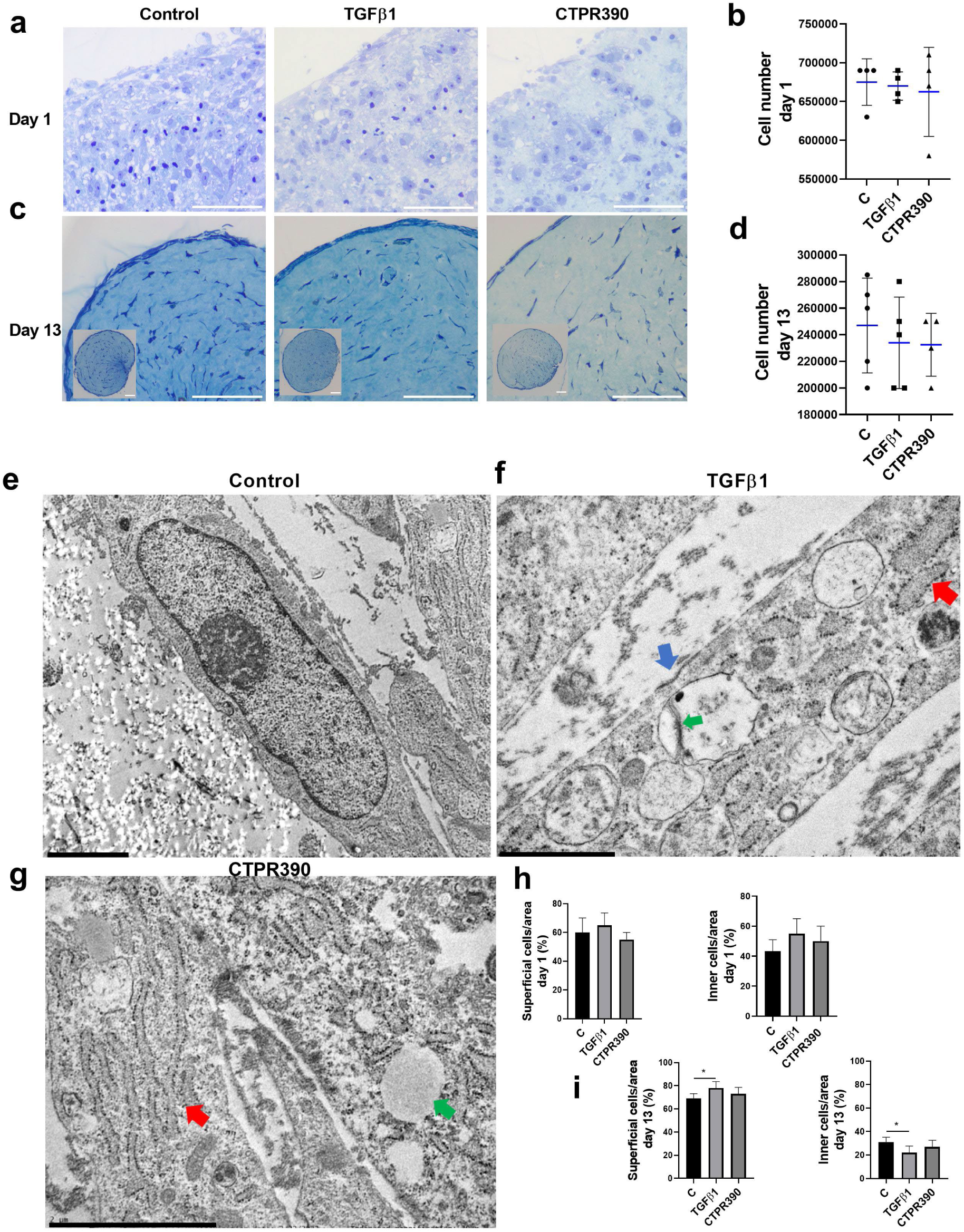
Characterization of distinct fibroblast subtypes in the ECT. **a, c**) Hematoxylin/eosin staining on day 1 (**a**) and day 13 (**c**) of cross-sections of Control, TGFβ1, and CTPR390 groups including insets of the whole cross section (**c**). The white bars of all panels indicated 50 μm, including inset panels showing the whole ECTs. **b, d**) Quantification of total cell numbers on day 1 (**b**) and day 13 (**d**) of Control, TGFβ1, and CTPR390 groups. Cell counts were performed in 3-5 biological samples per group including 3 technical replicates per sample. **e, f, g**) Electron microscopy images of superficial fibroblasts showing representative ultrastructural features of Control (**e**), TGFβ1 (**f**), and CTPR390 (**g**) ECT on day 13. **f**) The blue arrow marks a depression of the plasma membrane; the red arrow points to a sac of rough endoplasmic reticulum (RER); and the green arrow points to one of the vesicles with very electron-dense fibrous elements. **g**) The red arrow points to a normal-shaped RER sac; and the green arrow indicates a vacuole. Black scale bars indicated 2 μm (**e, g**) and 3 μm (**f**). **h, i**) Bar graphs showing the percentage of superficial and inner fibroblasts per area on day 1 (**h**) and day 13 (**i**); n = 3 biological samples per group in triplicate were measured. The p values (* p < 0.05) were determined using a one-way ANOVA with Tukey’s multiple comparison test.

A qualitative observation at the ultrastructural level of the fibroblasts of the TGFβ1-activated ECT revealed several key features. First, there was an electron-dense depression of the plasma membrane indicating active trafficking (blue arrow). Additionally, engorged sacs of rough endoplasmic reticulum (RER) were observed, suggesting a high level of protein transcription activity (red arrow). Finally, a vesicle containing very electron-dense fibrous elements was noted, which could likely be an exocytosis vesicle transporting fibers to the extracellular matrix (green arrow) (Figure 4f). On the other hand, the fibroblasts from CTPR390 ECT, as shown in the representative image of Figure 4g, presented normally shaped RER sacs (red arrow) comparable to the control (Figure 4e). However, they also showed the presence of vacuoles (green arrow), which did not appear under Control or TGFβ1 conditions. Nonetheless, this feature was not accompanied by any other signs of stress or apoptosis. Fibroblasts with a more rounded morphology, indicative of cells devoid of tension, were observed within the ECT in all groups (Supplementary Figure 3e-g) but not in the surface. We calculated the percentage of superficial fibroblasts relative to the total number of fibroblasts per area on day 1 and day 13. The results showed that the initial fibroblast distribution (day 1) was similar among groups and across the superficial and interior areas of the ECT (Figure 4h). On day 13, the total distribution of fibroblasts changed, showing a higher percentage of superficial fibroblasts in the TGFβ1-activated ECT compared to the Control group (Figure 4i), with a corresponding reduction in the percentage of inner fibroblasts compared to the Control group (Figure 4i). The CTPR390 group showed no significant differences compared to either the Control or TGFβ1 groups.

### CTPR390 diminished the production of structured collagen in fibrotic ECT

Collagen deposition during tissue remodeling is carefully regulated by local stresses and strains [36]. The ECT activated with TGFβ1 simulates tissue remodeling influenced by these local stress forces, as illustrated in Figure 1. Scanning electron microscopy revealed a tubular structure with layers of fibroblasts encasing the ECM on the tissue surface across all groups (Control, TGFβ1 and CTPR390) (Figure 5a). A detailed examination of the inner ECM in TGFβ1-activated ECTs revealed structures resembling cellular debris, indicative of poorer cell accommodation (Figure 5b). Further analysis revealed a significant reduction of elastin gene and protein, confirming the higher stiffness of the tissue (Supplementary Figure 4a-b). The TGFβ1 group displayed enhanced ECM activity, along with indicators of apoptosis and cell proliferation at both the gene and protein levels, which were not observed in the Control or CTPR390 ECTs (Supplementary Figures 4 c-h). Cross-sectional area (CSA) measurements of TGFβ1-activated ECTs showed a significant reduction compared to Control ECTs (****p < 0.0001) and CTPR390 ECTs (*p < 0.05) (Figure 5c). This reduction was attributed to matrix compaction by TGFβ1-activated fibroblasts. Fibrotic ECTs treated with CTPR390 showed partial protection from this compaction (*p < 0.05), corroborated by perimeter measurements (Figure 5d). A significant correlation between CSA and perimeter was observed in Control ECTs (****p < 0.0001) but disappeared with TGFβ1 treatment and was not restored by CTPR390 (Figure 5e). The polarized light technique differentiated between structured and unstructured collagen based on light refraction [37]. TGFβ1-activated ECTs displayed organized collagen fibers (light signal) compared to less organized collagen in Control ECTs (dark areas) (Figure 5f). Quantification revealed significantly more structured collagen in TGFβ1-activated ECTs compared to Control (**p < 0.001) or CTPR390-treated ECTs (**p < 0.001) (Figure 5g). To differentiate human ECM from bovine collagen of the ECT, we tested the species specificity of the human collagen Ia1 antibody. It showed no signal in bovine collagen, but positive labeling in ECT-surface collagen (Supplementary Figure 4i). TGFβ1-activated ECTs had a higher human collagen signal compared to Control and CTPR390 ECTs (Figure 5h), in a manner consistent with the marking by polarized light shown in Figure 5g. Multiphoton imaging, utilizing Second Harmonic Generation (SHG), identified areas of well-organized collagen (strong SHG signal) versus less organized collagen (weak or absent SHG signal). This technique confirmed high collagen organization in TGFβ1-activated ECTs and less organization in Control or CTPR390-treated ECTs (Figure 5i).

**Figure 5:**
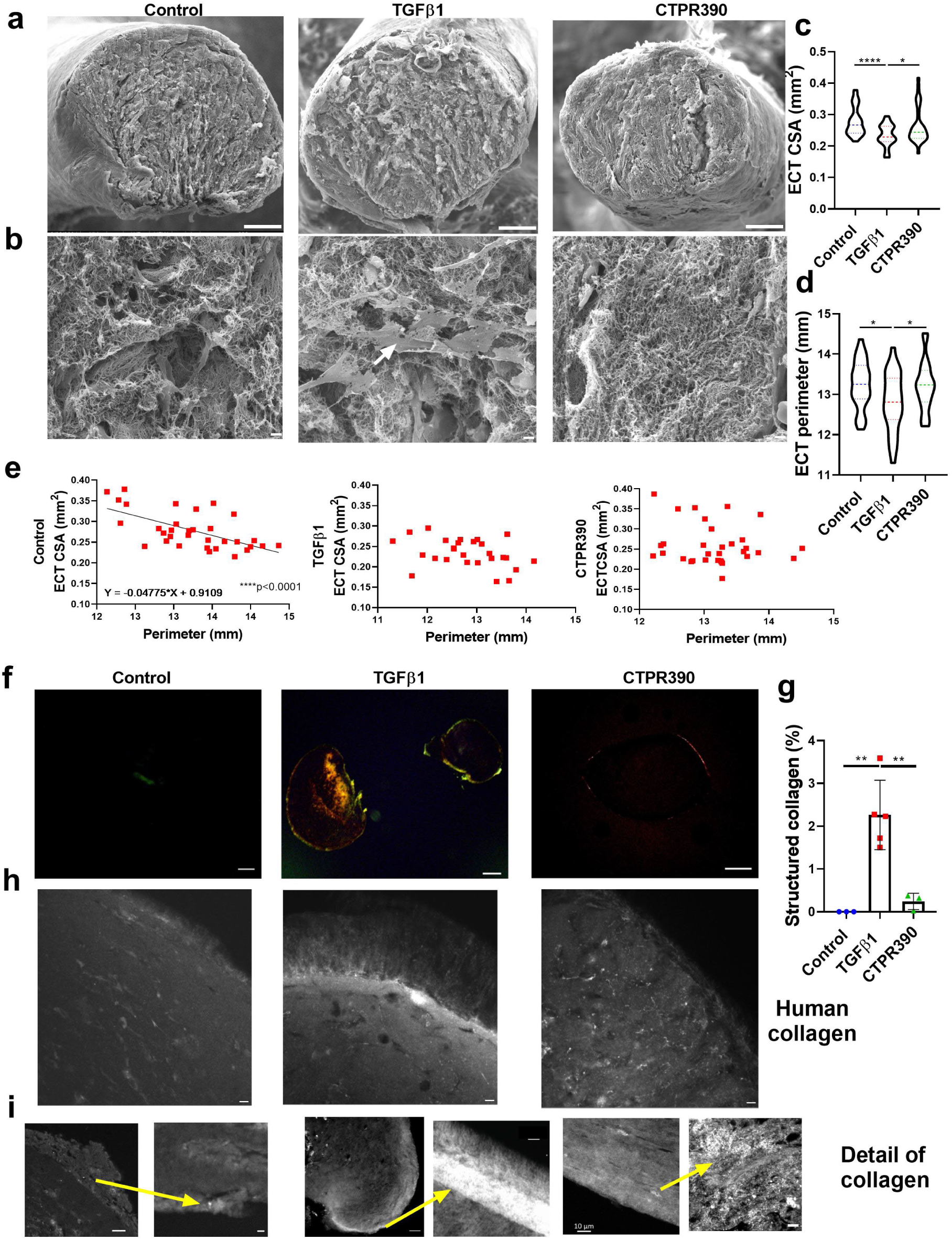
Localization of organized collagen produced by human fibroblasts on the surface of TGFβ1-activated ECT. **a**) Representative cross-sectional scanning microscopy images of Control, TGFβ1 and CTPR390 long arms of the ECTs. The white scale bars indicated 100 μm. **b**) Representative magnified scanning microscopy images of inner-sections of Control, TGFβ1 and CTPR390 ECTs. The white scale bars indicated 5 μm. **c, d**) Violin graph illustrating a significant reduction of the TGFβ1 ECT cross section area (CSA in mm^2^) (**c**), and a reduction of the TGFβ1 ECT perimeter (in mm) (**d**), indicating a higher degree of compaction in TGFβ1 samples in comparison to Control or CTPR390 ECTs (n=25-33 ECTs analyzed per group with 3 technical replicates per sample). **e**) Positive and significant correlation (****p<0.0001) between CSA and perimeter of Control ECTs. The other two groups (TGFβ1, and CTPR390) did not show a correlation between the CSA and their respective perimeters (n=25-33 ECTs analyzed per group with 3 technical replicates per sample). **f**) Representative images showing distinctively colored structured collagen at the periphery of the TGFβ1-activated ECT (central panel) under polarized light. In contrast, there was very low detection of structured collagen in the other two groups under study. The white scale bars indicated 50 μm. **g**) Bar graph showing the percentage of structured collagen assessed through polarized light; n = 4-5 ECT samples per group with 3 technical replicates per sample. The p values (*p < 0.05, **p < 0.005, ****p < 0.0001) were determined using a one-way ANOVA with Tukey’s multiple comparison test. **h**) Representative confocal imaging of the ECT surface detecting human collagen (COL I A1 antibody) showed a clear increase in collagen detection in TGFβ1 samples compared to Control or CTPR390 samples. The white scale bars indicated 10 μm. **i**) Second Harmonic Generation (SHG) signal showing the clear accumulation of collagen in TGFβ1 ECTs compared to Control or CTPR390 ECTs. Yellow arrows indicated the place of the augmentation. The white scale bars indicated 5 μm, except the fifth panel that indicated 10 μm.

### Fiber orientation preserved after CTPR390 treatment

Preferential angles and degree of fibers orientation (mainly collagen as shown in previous figures) were obtained from SAXS patterns and plotted as 2D maps (Figure 6a-c). We analyzed the level of collagen organization within each of the ECTs by analyzing the anisotropy of the scattering patterns (Figure 6, Supplementary Figures 5-7) [38]. The analysis conducted on each 20 x 20 μm² pixel image of the ECT surface revealed that the percentage of organized areas (those with a degree of orientation over 0.15) in the ECT was similar between Control and fibrotic ECT, with values of 91.7% in TGFβ1-activated ECT and 92.3% in Control ECT. However, a significant increase was observed in the organization level of fibers present in those extracellular matrix (ECM) areas in TGFβ1-activated ECT compared to Control ECT. This organization level was determined by the relative degree of orientation in TGFβ1-activated ECT, which was 0.56 ± 0.25 versus 0.52 ± 0.22 in Control ECT (Figure 6d). It is noteworthy that the fibers of TGFβ1-activated ECT appeared to avoid these high levels of organization after treatment with CTPR390 (Figure 6C-D). The significant decrease in both the extent of organized ECT areas and the degree of orientation of these areas in CTPR390-treated ECT was evident from the significantly lower values obtained for the percentage of structured area (80.9%) compared to TGFβ1-activated ECT, as well as the organization level of those areas (0.42 ± 0.26), which is significantly lower (****p < 0.0001) than the value for TGFβ1-activated ECT (0.56 ± 0.25) (Figure 6c-d). The significant decrease in both the extent of organized ECT areas and the degree of orientation of these areas in CTPR390-treated ECT was evident from the significantly lower values obtained for the percentage of structured area (80.9%) compared to TGFβ1-activated ECT, as well as the organization level of those areas (0.42 ± 0.26), which is significantly lower (****p<0.0001) than the value for TGFβ1-activated ECT (0.56 ± 0.25) (Figure 6d).

**Figure 6:**
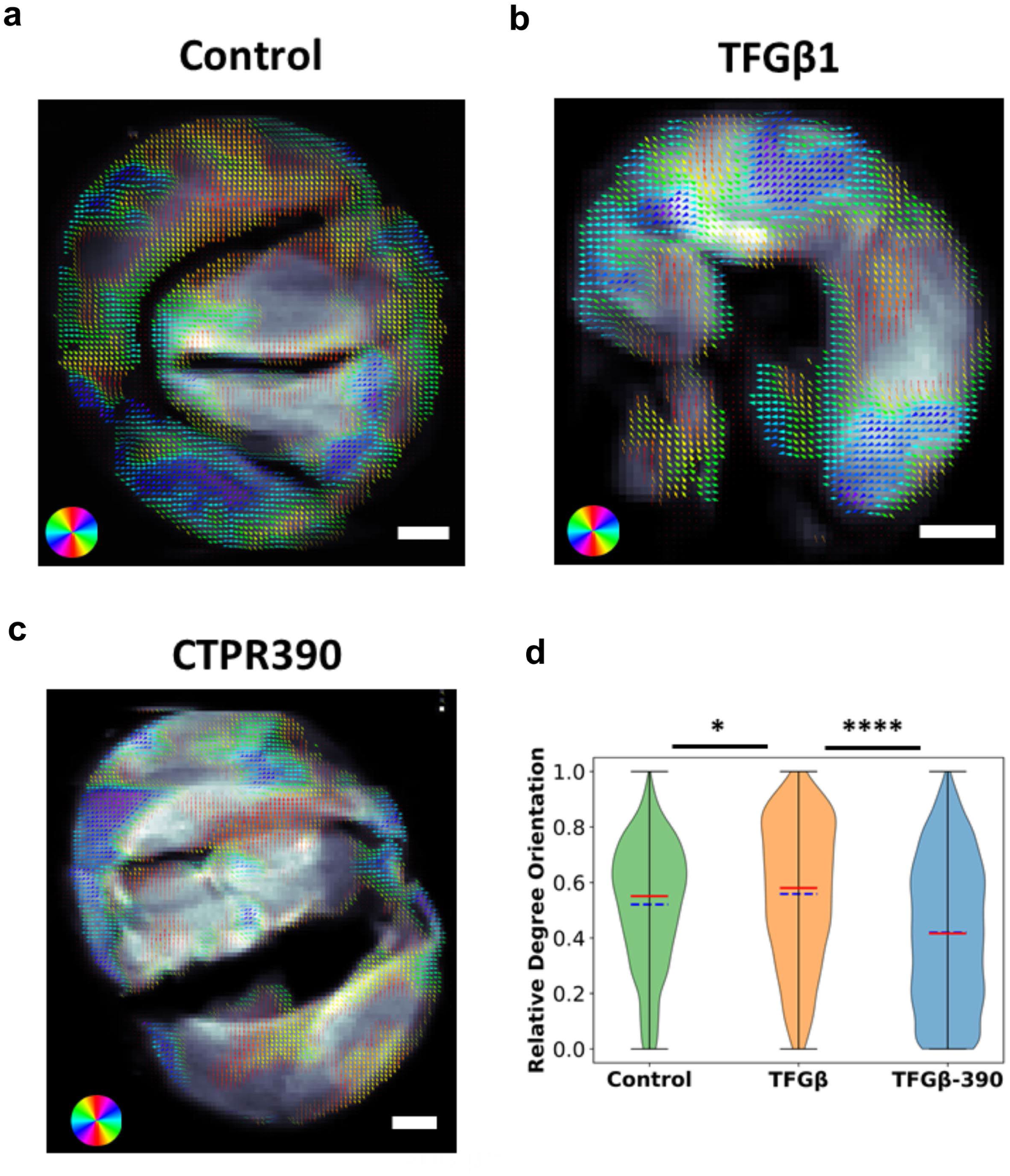
Differences in organization of collagen produced by human fibroblasts on the surface of TGFβ1-activated ECT. **a, b, c)** Maps of the preferential orientation and degree of orientation of collagen in control (**a**), TGFβ1(**b**), and CTPR390 (**c**) ECTs. The white scale bars indicated 200 μm. Preferential of orientation in each 20 x 20 µm^2^ pixel is indicated by the direction of the arrow in each of the pixels of the figure and the color coded discs. Degree of orientation in each 20 x 20 µm^2^ pixel is indicated by the thickness of the arrows in each of the pixels of the figure. Maps from individual ECTs are found expanded in Supplementary Figures 5-7. **d**) Violin graph illustrating significant changes in the degree of orientation of collagen in TGFβ1 (increase) and CTPR390 (decrease) ECTs. Graph was obtained considering the values from all of the pixels from each ECT analyzing one complete ECT per group including 3000-4000 pixels analyzed per ECT. The p values (*p < 0.05, ****p<0.0001) were determined using a one-way ANOVA with Tukey’s multiple comparison test.

## Discussion

In this study, we advanced preclinical human models to assess experimental drugs that had previously shown promising results in 2D cultures and *in vivo* models [19, 22]. Replicating the physical forces acting on the heart, including the ECM and fibroblasts, is crucial in a 3D model to capture their impact on morphology, biomechanics, and biochemical behavior. The adaptation used in this study accommodates such forces in both pathological and physiological scenarios, similar to approaches employed in other organs distinct from cardiac tissue [39–41]. Our ECT model, which mimics human cardiac scar tissue, helped identify the ultrastructural features associated with the anti-fibrotic treatment CTPR390. This treatment reduced pro-fibrotic markers in TGFβ1-activated human cardiac fibroblasts and altered fiber organization in the fibrotic ECT, making it resemble normal ECT. Previously, TGFβ1 activation induced pathological ECM changes that were reversed by CTPR390 in 2D cultures [31]. Moreover, the introduction of the hypertensive agent angiotensin II induced TGFβ1 activation, resulting in heart hypertrophy and fibrosis in a mouse model. This effect was successfully reversed through the administration of CTPR390 [19, 22]. Here, we show that CTPR390 restored biomechanical parameters in fibrotic ECT, addressing TGFβ1-induced changes. Treated ECTs had more variable fiber orientations and distributions, unlike the narrow alignment seen in fibrotic tissue. By segmenting fiber orientation, we detected alignment, providing new ways to study human cardiac fibrosis non-invasively. Tensile testing showed that CTPR390 prevented the increased stiffness of TGFβ1-activated ECT, returning it to control levels. Contractility measurements confirmed that treated ECTs exhibited biomechanical properties similar to controls. The biomechanical properties observed in treated ECT with CTPR390 were a direct outcome of its activity within fibroblasts. We demonstrated the effective penetration of the fluorescent CTPR390 into human fibroblasts within the tissue, showcasing its anti-fibrotic therapeutic properties. At the molecular level, these therapeutic capabilities, extensively validated in 2D and *in vivo* models [19, 22, 23], resulted in a reduction in the expression profile of cardiac fibrosis-associated genes in the human ECT. CTPR390 prevented the overexpression of key pro-fibrotic markers, including COL I A1, COL III A1, LOX, PDGFRA, and POSTN. Unlike in 2D cultures, protein regulation in ECT more accurately represented the pro- and anti-fibrotic effects of cardiac remodeling as it is the case for COL III, which has been implicated in altering the balance of collagen distribution during cardiac pathological remodeling to prevent cardiac dysfunction [42].

TGFβ1 activation heightened ECM production and increased fibroblast concentration in superficial tissue regions. Fibrotic human fibroblasts displayed greater activity, including filopodia and bleb formation, increased cell elongation, and larger cell areas and perimeters. CTPR390 treatment mitigated these pathological changes, reducing superficial fibroblast density and collagen formation, while maintaining the number of viable cells and preventing ECM accumulation in fibrotic tissue. These actions maintained an ECT architecture similar to control, thereby preventing the internal accumulation of additional ECM structures observed in fibrotic ECTs.

Ultrastructurally, CTPR390-treated superficial fibroblasts maintained their elongated shape and aligned with tissue forces. No morphological differences were seen in fibroblasts within the tissue core, regardless of treatment. This suggests that the fibrotic tissue reorganized itself, recruiting activated cells to regions driving tissue remodeling, leaving other areas unaffected. These actions could resemble the connective tissue regions affected by remodeling and those not affected by such remodeling in the heart, where resident fibroblasts become active or remain in a quiescent state depending on the needs. Fibroblasts of CTPR390-treated ECTs also displayed a return of the endoplasmic reticulum to baseline thickness, indicating a reduction in pro-fibrotic protein production. These changes aligned with normal myocardial conditions, where fibroblasts maintain heart homeostasis by contributing to ECM that distributes mechanical forces [43].

Our data demonstrated that ECT fibroblasts responded to pathological stimuli, similar to heart fibroblasts during post-inflammatory cardiac remodeling [44–47]. This phenomenon is observed in all types of cardiac fibrosis, including perivascular fibrosis, reactive fibrosis, and replacement fibrosis [48]. Cross-sections of fibrotic ECTs showed compaction and a less homogeneous ECM, which CTPR390 treatment prevented. In fibrotic areas, structured overexpressed collagen was observed, but CTPR390 reduced this to control levels. Additionally, CTPR390 treatment could not fully restore the correlation between compaction and perimeter values lost in fibrotic ECTs, and subtle differences, like vacuoles, were found in CTPR390-treated tissue. X-ray scattering patterns provided nanoscale data on collagen structured areas, revealing higher level of organization for collagen in fibrotic ECTs and less organized areas with lower level of structuration in treated and control tissues. Collagen orientation was linked to tissue stiffness, supporting findings from PS-OCT. The structural characterization validated collagen’s biomechanical properties across control, fibrotic, and anti-fibrotic ECTs.

In conclusion, our findings underscore the intricate relationship between ECM and fibroblasts in tissue biomechanics, offering insights for anti-fibrotic therapies and diagnosis.

## Online Materials and Methods

### Cell culture measurements of primary human cardiac fibroblasts

We obtained adult human left ventricular cardiac fibroblasts from a male Caucasian donor through Promocell (Lot 437Z012.4). The cells were cultivated as a monolayer at 37 °C under 5% CO_2_ in humidified incubators, using FGM-3 growth medium (PromoCell) supplemented with 10% Fetal Calf Serum, Basic Fibroblast Growth Factor 1 ng / mL and Insulin 5 μg / mL, without adding antibiotics, following the manufacturer’s guidelines.

### Preparation of human engineered connective tissue

The adult human left ventricular cardiac fibroblasts were expanded, and assayed for ECT production were conducted between passages 3-4. The ECT generation was performed as described before [12]. In brief, all required materials were pre-chilled, and all following steps were performed on ice. First, 0.3 mg/ECT bovine collagen type I (Collagen solutions LLC) was mixed with 2 × DMEM and neutralized with 0.2 M NaOH. Then, resuspended 7.5 × 10^5^ human cardiac fibroblast per ECT in FGM-3 media were added and thoroughly mixed. The final volume per ECT was 180 µL. The cell-collagen mixture was pipetted in 48-well mold plates containing two flexible poles containing a specific material named TM5MED (shore value A46, bending stiffness 1.5 mN/mm) (myrPlates myriaMed) [16, 49] and let condensate for 1 h at 37 °C in a cell incubator. The ECT were cultured for 13 days, and the medium, including the additives, was changed every second day. The concentration of TGFβ1 used in this study (5 ng/mL) was consistent with the one previously employed in this model to induce fibrosis [16]. The optimal anti-fibrotic concentration of CTPR390 was previously established to be 1 μM based on the extrapolation of titration assays conducted in primary fibroblasts [19].

### ProtCTPR390 expression and purification

CTPR8(C390-C416His-E2C) (ProtCTPR390) was designed as a multifunctional protein scaffold. The chimera protein includes several modules: a wildtype module with the glutamate residue at position 2 (E2) mutated to cysteine to allow fluorophore conjugation through the highly selective and straightforward thiol-maleimide chemistry; a CTPR390 module for Hsp90 binding functionality, and a CTPR416His-E2C module with four tandem repeated CTPR units containing histidines at positions 2, 5, 9, and 13 of each repeat, serving as a metal coordination domain to stabilize the formation of AuNC. This module presents an additional point mutation to insert a single Cysteine for conjugation of selected fluorescence dyes (E2C). The plasmid encoding the CTPR8(C390-C416His-E2C) gene (ProtCTPR390) was transformed into Escherichia coli C41 (DE3). The protein of interest was expressed by culturing the bacteria under agitation at 37 °C, and induced by the addition of 1 mM isopropyl β-d-thiogalactoside (IPTG) when they reached an optical density of 0.6. Then, the cultures were grown overnight at 20 °C. Cells were pelleted and resuspended in 50 mM Tris, 500 mM NaCl, pH 8.0. The cells were lysed through freeze-thaw cycles and lysed by sonication and centrifuged for 1 h at 10000 rpm. The overexpressed His-tagged CTPR390 protein was purified by affinity chromatography using a 5 mL HisTrap Q column (GE Healthcare). The His6-tag was then cleaved by Tobacco Etch Virus (TEV) protease overnight at room temperature, and then purified by size-exclusion chromatography. The protein concentration was determined by measuring the absorbance at 280 nm, using extinction coefficient calculated from the amino acid composition. Purity and size of the protein were confirmed by sodium dodecyl sulfate - acrylamide electrophoresis gel (SDS-PAGE), and by Matrix-Assisted Laser Desorption/Ionization-Time of Flight (MALDI-TOF) using an Applied Biosystems Voyager Elite MALDI-TOF mass spectrometer with delayed extraction (Applied Biosystems, Framingham, MA, USA). The protein secondary structure was verified by circular dichroism using a Jasco J-815 (JASCO Corporation, Tokyo, Japan). CD spectra were obtained at 2.5 μM of protein using a cuvette with 1 mm path length.

### Synthesis of protein-stabilized gold nanoclusters (CTPRAu)

AuNC were stabilized on ProtCTPR390 protein through biomineralization, by optimizing a protocol previously described before [25, 49]. Initially the protein buffer was changed to 150 mM of NaCl, 50 mM phosphate buffer at pH 10 using a PD-10 desalting column. Then, 1 mL of protein at 40 μM was incubated with 80 equivalents of Au per protein (21.4 μL of HAuCl4 at 150 mM) at 50 °C for 30 min. Then, the protein was placed for 10 min on ice, followed by the addition of 8000 equivalents (213 μL at 1.5 M) of sodium ascorbate and incubation at 50 °C for 72 h. After the synthesis, CTPR390-NC complex was centrifuged at 16,400 rpm for 1 h and subsequently concentrated using an Amicon filter with pore size of 10 kDa. Finally, the protein-nanomaterial hybrid was purified using a PD-10 desalting column and the buffer changed to PBS for further storage and use [25].

### Conjugation of CTPR390-AuNC with Alexa Fluor 488

After the synthesis of CTPR390-AuNC, the cysteine in the protein scaffold was conjugated with Maleimide C5 Alexa Fluor™ 488 (Thermo Fischer). For this, CTPR390-AuNC in PBS pH 7.4 were incubated with 10 equivalents of dithiothreitol (DTT) for 30 minutes at 4 °C. Afterwards, the DTT was removed using a PD-10 column and CTPR390-AuNC incubated with 13 equivalents of Maleimide C5 Alexa Fluor™ 488. The reaction was incubated for 4 h at 25 °C. After the conjugation reaction the solution was concentrated, and the excess of dye removed using a PD-10 desalting column. All the samples were sterilized using a 0.22 μm syringe filter under sterile environment and stored in the fridge until further use. The protein concentration in the CTPR390-AuNC and CTPR390-AuNC-488 samples was determined by Pierce™ BCA Protein Assay Kits (Thermo Scientific™) using as calibration line the same protein scaffold used for the synthesis of the hybrid nanomaterials (CTPR390). The amount of gold was determined by inductively coupled plasma mass spectrometry (ICP-MS, iCAP-Q ICP-MS, Thermo Scientific, Bremen, Germany) of samples previously digested with aqua regia. In summary, the CTPR390 used in this study contained its characteristic Hsp90 binding module fused to a histidine-based gold coordination domain and was mutated with a single cysteine for the conjugation with maleimide-Alexa 488 fluorophore. This protein was subsequently used to stabilize the formation of gold nanocluster (CTPR390-AuNC) and conjugated with Alexa 488 (CTPR390-AuNC-488).

### CTPR390 fluorescence

The ProtCTPR390, CTPR390-AuNC, and CTPR390-AuNC-488 (CTPR390-AuNC-488 is named as CTPR390 in the main text) fluorescence was evaluated using a BioTek Synergy H1 hybrid microplate reader in a black 384-well plate. The fluorescence spectrum was recorded in RFU (Relative Fluorescence Units) against wavelength (nm) plots through a monochromator (1 second of integration time), measuring by the top, reading height of 7 mm, and gain of 100. The emission spectra were obtained by exciting the sample at 460 nm and recording the emission from 500 to 800 nm, or exciting the sample at 360 nm and recording the emission from 400 to 800 nm with a step of 2 nm.

### ECT compaction analysis by cross-sectional area measurement

Tissue compaction, which starts after collagen gelation, involves cell-driven matrix compression and remodeling. This is measured by the cross-sectional area (CSA) and volume of the tissues. On day 13, tissues were transferred to a DPBS multi-well plate, and top and side view images of the engineered cardiac tissues (ECTs) were captured using a Lumar.V12 stereomicroscope.

Tissue diameters were analyzed at a minimum of 8 positions per imaging plane using ImageJ. Mean diameters from top (dt) and side (ds) views were calculated, and CSA was determined using the equation: CSA = π × (dt/2) × (ds/2). The volume was calculated by multiplying the averaged CSA by the full length, measured from the top view using ImageJ.

### Stress-strain analysis of destructive tensile strength measurements. Pole deflection analysis

To evaluate the mechanical properties of engineered connective tissues (ECT), destructive tensile strength measurements were performed using an RSA-G2 rheometer. ECT samples were mounted on custom hooks, produced via 3D printing in-house, in a 37°C organ bath filled with DPBS and stretched uniaxially at a constant rate until rupture. ECTs were stretched at 0.03 mm/s. Force values were divided by cross-sectional area (CSA) to obtain stress values, which were then plotted against the gap between hooks. The stress-strain curves were analyzed to identify the toe region, elastic region, and plastic region.

To calculate the given parameters (Young’s modulus, strain to failure point) from the destructive tensile strength measurements, the measured force values (mN) were divided by the determined CSA (mm^2^) to obtain stress values (kPa) and these were plotted against the gap in mm (distance between the upper and lower hook) in GraphPad Prism. The obtained curves typically displayed the following regions: An initial region in which only background stress was measured, a short toe region, an elastic region in which the stress versus gap values were linear, a flattened plastic region, and finally a region of failure which was characterized by a sudden drop in stress due to the rupture of the tissues. To calculate the strain values, the beginning of the toe region was identified, and the corresponding gap length was considered as L0. The strain equation (Ltotal-L0)/L0 was used to transform the gap values, and the stress values at the beginning of the toe region were used for background subtraction. The slope of the elastic region was determined by linear regression and is presented in this work as Young’s modulus. The strain to failure point, characterized by the sudden drop in stress, was identified manually. Due to the material-dependent variations in the absolute values, we compared only conditions which had been tested in parallel. Maximum stress, ultimate strain, and resilience were calculated using area under the curve (AUC) analysis. All assays were performed with n= 12 ECT per group.

ECT pole deflection were imaged daily over a 13-day period using a Nikon D90 camera with a Nikon AF-S NIKKOR 18-105mm lens. UVA light was used to enhance the visibility of the poles, allowing for precise measurement of their distance on the plate under sterile conditions. This setup ensured non-invasive, high-resolution imaging throughout the experiment. Images were analyzed with ImageJ software to measure the distance between the poles, which indicated the degree of pole defletion. Changes in pole distance was quantified relative to the initial measurement on day zero, providing a reliable measure of tissue biomechanical forces over the 13 days.

### Enzymatic digestion of ECT to disaggregate primary human fibroblasts

For each experimental condition (Control, TGFβ1, CTPR390, n=5), ECT organoids were digested using a two-step enzymatic process. Initially, organoids were treated with a 2 mg/mL Collagenase I solution for 3 h at 37°C. This was followed by a 30-minute incubation with Accutase (detachment solution of proteolytic and collagenolytic enzymes) at 37°C. After each treatment, the resulting solution was collected and kept on ice. DPBS with 5% FBS was added to inactivate the enzymes. The combined cell solution was then centrifuged, and the cell pellet resuspended in culture medium. The cells were subsequently used for viability assays, cell counting, or prepared for microscopic observation.

### Fluorescence Live Cell and ECT Imaging

On day 13, control ECT, TGFβ1-activated ECT and TGFβ1-activated ECT and treated with CTPR390 were assessed for fluorescence. Excitation and emission were set at 488 nm and 495 nm, respectively to detect Alexa 488 fluorophore conjugated to CTPR390, Nikon Eclipse Ti2 microscope was employed. On day 13, the human left ventricular cardiac fibroblasts embedded in ECT were dissociated by incubating them in 2 mg/mL collagenase I in calcium-containing PBS in the presence of 20% FCS for 2 h at 37 °C. Subsequently, the tissues were washed with PBS and further incubated in Accutase (Millipore), 0.0125% Trypsin, and 20 μg/mL DNAse (Calbiochem) for 30 min at 37 °C. The cells were mechanically separated, and the enzymatic activity was halted by transferring them into PBS containing 5% FBS. After centrifugation for 10 minutes at 100 g and 4 °C, the cells were resuspended in FGM-3 growth medium (Lonza) supplemented with 10% Fetal Bovine Serum (FBS, Gibco), 100 U / mL penicillin, and 100 μg/mL streptomycin (Gibco). The cells were either added to PBS containing 5% FBS and counted using a CASY TT system (Roche), or they were seeded in 35mm microscopy dishes (Ibidi µ-Dish 35 mm high Glass Bottom 81158) and incubated for 2 h at 37 °C under 5% CO_2_ in a humidified incubator. After washing, the medium was replaced with Leibovitz microscopy medium (Gibco™ 21083027). Inverted transmitted-light microscope, Primovert 491206-0001-000 Zeiss was used for the *in vivo* ECT confocal images. Fluorescence belonging to the CTPR390 incorporated to the fresh ECT was detected with Orca flash 4 as a camera. A 60X oil objective with an opening of 1.4 mm was used. To detect CTPR390 –Alexa 488 fluorophore, excitation and emission were set at 488 nm and 495 nm, respectively. Objective 4x / Plan-Achromat 4x/0.10 for Primo Objective 10×/ Plan-Achromat 10×/0.25 for Primo. Nikon Eclipse Ti2 was employed for 2D cell live image acquisition.

### Polarization-sensitive optical coherence tomography

Polarization-sensitive optical coherence tomography distinguishes itself as a method with significant potential for offering a comprehensive depiction of the morphological architecture of tissues. Samples were measured with the TEL221-PS commercial system (Thorlabs Inc.), with 13 μm/pixel lateral and 5.5/n μm/pixel depth resolution. For the purpose of this work, n = 1.38 was used as the refractive index of all samples. Samples were imaged in a single sweep in approximately one minute each. The system illuminates the sample with circularly polarized light to excite all orientations of the sample’s fibers equally. Afterwards, the parallel (⟨*E*^2^_*x*_⟩) and perpendicular (⟨*E*^2^_*y*_⟩) components of the electric field are captured, and the Stokes parameters (*s⃗* = (*I*, *Q*, *U*, *V*)) are calculated (*I* = ⟨*E*^2^_*x*_⟩ + ⟨*E*^2^_*y*_⟩, *Q* = ⟨*E*^2^_*x*_⟩ − ⟨*E*^2^_*y*_⟩, *U* = 2⟨*E*_*x*_ *E*_*y*_ *cos*(δ)⟩, *V* = 2⟨*E*_*x*_ *E*_*y*_ *sin*(δ)) at all points of the three-dimensional PS-OCT cube. The angle of polarization is then calculated as 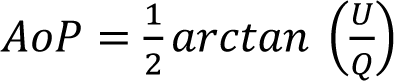. To find the attenuation coefficient (μ_*z*_), the Beer-Lambert law was applied to fit the PS-OCT signals to *I* = *I*_*z*=0_*e*^−2μ*_z_z*^, where *I*_*z*=0_ represents the intensity (Stokes I) at the surface of the sample. The surface profile was obtained as the maximum intensity along depth for each xy position (*max*(*I*(*x*, *y*, *z*)|_*z*_). Kernel Density Estimation (KDE) was used to find the 2D distributions of the samples in the U-μ_*z*_ plane. The distributions were then bound to their 1σ areas to find the overlap between samples by discretizing the space with a uniform 100 x 100 grid. Total areas analyzed excluded poles contact area previously described as intrinsic fibrotic areas [16]. This measurements corresponded to the evolution along the depth (z) of the xy-averaged Stokes parameter (U) that characterized differences in tissue organization. The steady rate of change along depth (0-100 µm) corresponding to a linear relationship between U and z indicates some level of fiber uniformity (overall change below - 10% for all samples), with steeper slopes being markers of higher fiber alignment between different layers of the sample.

### Immunochemistry of ECT. TUNEL assays

ECT previously fixed in PFD 4% and embedded in paraffin were permeabilized 30 min with 0.05% Triton-X in PBS. Immunochemical detection was carried out by incubating overnight at 4 °C with the following anti human antibodies: anti-FAP antibody (1:100 dilution, PA5-51057), anti COL I A1 antibody (1:100 dilution, ab34710), anti COL III A1 antibody (1:100 dilution, ab184993), FAP antibody (1:100 dilution, PA5-51057), anti-αSMA antibody (1:100 dilution, A5228), anti-Ki67 MIB-1 (1:1000 dilution, Dako M7240), anti-elastin (1:1000 dilution, A-8: sc-374638). The samples were washed with 0.05% Tween 20 in PBS, incubated for 1 h at room temperature with a secondary antibody (anti-human Cy5), washed in PBS. Fixated ECT were left in PBS at 4 °C for the subsequent confocal imaging. The same protocol was followed for the apoptosis assays using In Situ Cell Death Detection Kit (12156792910), TMR red. Terminal deoxynucleotidyl transferase-mediated TMR dUTP nick-end labeling (TUNEL) staining. An in situ cell death detection kit (Roche Diagnostics, Penzberg, Germany) was employed to detect apoptotic cells. ECT vibratome sections (100 μm) were rinsed in PBS (3 × 10 min) and permeabilized at room temperature for 30 min with 0.5% Triton X-100 diluted in PBS. ECT sections were incubated at 37 °C with the TUNEL reaction mixture containing TMR-conjugated dUTP, terminal deoxynucleotidyl transferase and nucleotide mixture for 1h in a wet chamber. The samples were then washed with PBS and mounted with antifading solution. TUNEL staining was also combined with DAPI, mounted with vectashield and confocal images were processed with ImageJ.

### *In vitro* treatment of ECT with TGFβ1 and CTPR390

ECT acceptor generated as explain above and incubated at 37 °C with 5% CO_2_. After 24h, the correct compaction to the poles was reached. At this point, 5 ng/mL of TGFβ1 and 1 μM of CTPR390 final concentration were added to the media. New media including fresh TGFβ1 for TGFβ1-activated ECT, and fresh TGFβ1 (5 ng/mL) with CTPR390 (1 μM) was added every 4 days to TGFβ1-activated ECT treated with CTPR390 group.

### Quantitative PCR analysis of cell samples

The total RNA from fibroblasts embedded into ECT was obtained with TRIzol® reagent (Gibco BRL, Grand Island, NY). Complementary DNA was prepared from 0.5 μg total RNA by random priming using a first-strand cDNA synthesis kit (Promega Corp). Primers sequence for the human genes using Sybr green QPCR were the following: COL IA1 forward: CCAGCAGATCGAGAACATCC; reverse: CAGAGTGGCACATCTTGAGG. COL IIIA1 forward: GACCAAAAGGTGATGCTGGC; reverse: ACCGTTAGCTCCTGGTTTCC. LOX I forward: CAAGGGACATCAGATTTCTTACC; reverse: CCATACTGTGGTAATGTTGATGAC. CTGF forward: GAGAGTCCTTCCAGAGCAGC; reverse: CATAGTTGGGTCTGGGCCAA. PDGFRA forward: GCTCTTTACTCCATGTGTGGGA; reverse: ATTAGGCTCAGCCCTGTGAGA. TCF21 forward: AACGACAAATACGAGAACGGGT; reverse: CTCCAGGTACCAAACTCCAAGG. FAP forward: GCTTTGAAAAATATCCAGCTGCC; reverse: ACCACCATACACTTGAATTAGCA. FN forward: ACAACACCGAGGTGACTGAGAC. reverse: GGACACAACGATGGTTCCTGAG. ACTA2 forward: AGAACATGGCATCATCACCA. reverse: GCGTCCAGAGGCATAGAGAG. POSNT forward: CCCTTGGAAGAGACGGTCAC. reverse: CTCAAAGACTGCTCCTCCCA. Elastin forward: TTCCCCGCAGTTACCTTTCC. reverse: CTAAGCCACCAACTCCTGGG. Ki67 forward: ATTTGCTTCTGGCCTTCCCC. reverse: CCAAACAAGCAGGTGCTGAG. CCND1 forward: CTGATTGGACAGGCATGGGT. reverse: GTGCCTGGAAGTCAACGGTA. FGF2 forward: CCCCAGAAAACCCGAGCGA. reverse: TTCACGGATGGGTGTCTCCG. PDGFRB forward: CAGCTCTGGCCCTCAAAGG. reverse: GAACGAAGGTGCTGGAGACA. BAX forward: AGGGGCCCTTTTGCTTCAG. reverse: TGTCCAGCCCATGATGGTTC. BCL2 forward: GAAGGTTTCCTCGTCCCTGG. reverse: GAAGACCCTGAAGGACAGCC. P53 forward: AGTCTAGAGCCACCGTCCAG. reverse: TCCGGGGACAGCATCAAATC. P21 forward: AGTCAGTTCCTTGTGGAGCC. reverse: CATTAGCGCATCACAGTCGC. GREM1 forward: TAAGCAGACCATCCACGAGG. reverse: GGCAGTTGAGTGTGACCATC. FSP1 forward: CCCTGGATGTGATGGTGTCC. reverse: CGATGCAGGACAGGAAGACA. HSP90AA1 forward: GGTCCTGTGCGGTCACTTAG reverse: TATCTGCACCAGCCTGCAAA HSP90AB1 forward: GGGGGTGTGTGGTAAGCAA reverse: CCATCACAGCCCCCATAGTT.

The target mRNA expression levels were normalized to GAPDH levels. GAPDH forward: TGCACCACCAACTGCTTAGC; reverse: GGCATGGACTGTGGTCATGAG. Relative quantization was expressed as fold-induction compared with controls in triplicate of three independent experiments.

### Confocal fluorescence microscopy of ECT cross-sections

Leica SP5 confocal, 63X oil objective, 1.4 mm aperture and 2.5 zoom was used. Fluorescence of ECT cross-sections was taken using 49% intensity, CTPR390 (excitation: 390 nm, emission: 488 nm), DAPI channel, laser 405 nm, intensity 27%, excitation filter 415 nm, emission filter 500 nm. Seven z-slices were acquired with a step size of 1.13 μm. The images were analyzed using ImageJ software. A median filter was applied to each channel and a maximum intensity z-projection was performed for each channel separately.

### Transmission Electron Microscopy

Control, TGFβ and TGFβ CTPR390-488 treated ECTs were examined for conventional ultrastructural analysis. ECTs were fixed with 1% paraformaldehyde and 1% glutaraldehyde in 0.1 M phosphate buffer, pH 7.4. Then they were cut in half to get material for TEM and Scanning Electron Microscopy (SEM) analysis. Continuing with the TEM protocol, the ECTs were rinsed in 0.1 M phosphate buffer, postfixed in 2% osmium tetroxide, dehydrated in ethanol, and embedded in araldite (Durcupan, Fluka, Switzerland). Semi thin sections were used not only as a quality proof to proceed to ultrathin sections but also for quantification, in that case, ultrathin sections were stained with 1% toluidine blue and observed and digitalized with a conventional light microscope (Leica DM1000) and analysed with ImageJ software [50]. All the analyses were carried out using GraphPad software for Windows. Ultrathin sections were transferred to copper grids and stained with uranyl acetate and lead citrate. Transmission Electron Microscopy (TEM) images of cross-section of ECT were recorded at an accelerating voltage of 80 kV and with magnifications ranging from 6000× to 29,000× using a GatanUltraScan 100 slow-scan CCD camera and software Gatan.

### Scanning Electron Microscopy

The sectioned ECTs were fixed in 3% glutaraldehyde, dehydrated with a graded ethanol series, dried by the critical point method, coated with gold in a Fine coat ion sputter JFC-1100 226 (JEOL, Ltd), and observed with an Inspect S microscope (FEI Company) working at 25 kV.

### Cellular area, perimeter and layers quantification

To measure the area and perimeter of cells extracted from engineered connective tissues (ECT), the following procedure was used. First, ECT samples were dissociated to obtain individual cells, which were then purified and replated for 24 h. After this incubation period, cell images were captured using an Olympus IX81 fluorescence microscope with a 10× UPlanFLN objective. These images were subsequently analyzed with ImageJ software to measure the perimeter and area of the cells. This method provided accurate measurements of cell morphology and spatial organization.

For the measurement of the number of cell layers, various areas within the longitudinal semi-thin sections of the three ECT groups under study (Control, TGFβ1, CTPR390) were analyzed. The superficial cell count was performed covering areas of 4,000 μm², while the interior cell count was conducted in areas of 16,000 μm². The relativization of the percentages of superficial cells and interior cells of the ECT was done in relation to each of the analyzed areas.

### Cell viability and counting method

Cell parameter values for number, viability, and diameter were obtained using the Casy TTS system, which utilizes three-dimensional counting, electrical current exclusion (ECE), and pulse area analysis for accurate measurements. During measurement, cells in either growth media or 5% FBS-containing DPBS were kept on ice. CASY accurately determines cell viability without interference from staining or focusing. Viable cells are identified by their intact cell membranes, which act as a barrier to current, while dead cells are detected by their nuclei due to the lack of membrane resistance.

### Polarized light imaging of Picrosirius Red stained ECT samples

Collagen fibers, when organized in a more structured and aligned manner, exhibit birefringence or the ability of a material to refract light in two different directions. The birefringence given by the polarized light passing through structured collagen of ECTs enabled us to distinguish between regions of the ECT characterized by organized collagen and those with less organized or unstructured collagen. Collagen content in ECT constructs (n = 2/condition: Control, TGFβ1, CTPR390) was quantified using a Picrosirius Red staining protocol. Briefly, ECT samples were fixed in 4% formaldehyde, paraffin-embedded, and sectioned into thin 0.5μm cross-sections. Following deparaffinization and rehydration through a graded series of xylene and ethanol washes, sections were stained with 1% Picrosirius red solution (Sirius Red in picric acid). Excess stain was removed by washing, and sections were dehydrated and mounted for subsequent analysis of collagen fiber cross-sectional area. Collagen fibers will be observed as birefringent, i.e., with bright colors ranging from red to yellow-green, depending on their orientation in the field of view. This allowed for the measurement of the cross-sectional areas of the ECT samples. Polarized light images were acquired using a Zeiss Cell Axio Observer microscope. Transmission imaging was performed with an EC Plan Neofluar 10x air objective (0.3 numerical aperture) and a Zeiss 305 Axiocam, utilizing both brightfield and polarization modes (0° polarizer, 90° analyzer). The total area of ECT was calculated through transmission image processing, and collagen quantification was achieved by analyzing the positive areas using the open-source software Fiji [50].

### Multiphoton microscopy

The multiphoton imaging technique utilizes nonlinear optical effects to visualize collagen structures in biological tissues. Specifically, it relies on two-photon excitation, where two photons of lower energy combine to excite a fluorophore. In the case of collagen, Second Harmonic Generation (SHG) is the predominant signal observed. Structured collagen, such as that found in organized and aligned fibers, produces a strong and specific SHG signal. This is because the nonlinear optical process is enhanced in highly ordered structures. To detect structured collagen, the Second Harmonic Generation (SHG) nonlinear optical process was employed (n = 2 ECT per group) using a Zeiss LSM 880 NLO multiphoton laser microscope coupled with a Mai Tai Deep See multiphoton laser. Imaging was performed with a C Apochromat 40x water objective (1.20 numerical aperture). Big2 detectors (Non-Descanned Detectors) were used for image acquisition in the 500-550 nm range, with excitation at 780 nm at 1% power. The image pixel size was set to 0.10 micrometers, with 8-bit resolution.

### SAXS imaging

Synchrotron Small-Angle X-ray Scattering (SAXS) imaging was collected at the SAXSMAT [38] beamline in PETRAIII from three groups of ECT cultivated for 13 days: Control, TGFβ1 (to induce fibrosis) and CTPR390 to prevent fibrosis. ECTs were extracted from the cultured plate and transferred to the planchettes for their cryopreservation in a 20% BSA freezing media. Then, they were cryofixed in an EM HPM100 High Pressure Freezing System and freeze died following temperature and pressure gradients with a single chamber method in a Chris Alpha 2-4 LSCplus. This generated ECT samples around 3 mm in diameter and 0.3 mm thick, ready for their study at room temperature. SAXS studies were performed at the P62 SAXSMAT beamline (PETRA III, Hamburg, Germany) [51] with a 12 keV beam focused at 21 x 22 µm^2^, and using an Eiger 2 X 9 M detector (in an evacuated flight tube) at a distance of 4.7 m. An ionization chamber before the sample and a beamstop in front of the SAXS detector allowed to monitor primary and transmitted beam intensities (for normalisation purposes). The ECTs were wrapped and immobilised using Kapton foil, and positioned on a goniometer inside a chamber filled with He (to minimize air background scattering). SAXS maps were acquired from areas between 4-6 mm^2^ that were continuously raster scanned with a virtual spot size of 20 x 20 µm^2^ and 0.2 s dwell time per virtual point of the YZ projections. Each 2D SAXS pattern was transformed to show scattering values between 0° and 360°. Afterwards, the one-dimensional SAXS profiles were corrected by self-attenuation, effective exposure time per virtual point, normalized by the incoming flux, and the background signal. The 5^th^ reflection order of collagen (q ≈ 0.48 nm^−1^) was used to reconstruct the collagen intensity, rank-1 tensor maps and degree of orientation as previously described [38]. The degree of orientation values for each ECTs were normalized to the highest alignment signal among the three ECTs for clarity. Any pixel with a degree of orientation below 0.15 was considered as an area of relative disoriented collagen. To avoid unwanted beam damage during acquisition, SAXS imaging were performed using dry samples. Yet, due to their size, the different ECTs studied needed to be cut and folded inside the aluminum capsules used for cryofixation under high pressure, and freeze dried before analysis. Unfortunately, this hampered recognition of the different sections from the ETCs within the images generated. Initially, the intensity of collagen alignment in each 20 x 20 μm² pixel of the ECTs was normalized to the highest alignment signal among the three ECTs, to obtain a value of relative degree of orientation (between 0 and 1). Any pixel with a relative degree of orientation below 0.15 was considered as an area of relative disoriented collagen.

## Statistical analysis

Data presented in this study were expressed as means ± SEM for n = 3-24 ECT per group depending on the assay performed. P values (* p < 0.05, **p < 0.005, ***p<0.0005, ****p < 0.0001) were determined using 1-way, and 2-way ANOVA with Tukey’s multiple comparison test.

## Acknowledgements

A.V.V. This work was partially supported by Agencia Estatal de Investigación (AEI/MCI/10.13039/501100011033), Spain PID2021-125702OB-I00). Fomento de la transferencia de conocimiento en la Comunidad Autónoma de Cantabria SUBVTC-2023-0006 T. A.L.C. acknowledges support by the Agencia Estatal de Investigación Grant PDC2021-120957-I00-NanoIVD and PDC2022-133345-I00-ProIMAGE funded by MCIN/AEI/ 10.13039/501100011033 and by the “European Union NextGenerationEU/PRTR” and Grant PID2022-137977OB-I00-ProTher funded by MCIN/AEI/ 10.13039/501100011033. C.S.C. thanks Gipuzkoa Foru Aldundia (Gipuzkoa Fellows program) Grant number 2019-FELL-000018-01/62/2019 for financial support. G.G. thanks the financial support of “la Caixa” Foundation (ID100010434, fellowship: LCF/BQ/DI20/11780020). This work was performed under the Maria de Maeztu Units of Excellence and Severo Ochoa Centres Programs from the Spanish State Research Agency – Grants No. MDM-2017-0720 (CIC biomaGUNE) and CEX2018-000867-S (DIPC). O.M.C. and V.M. thank the AEI for the infrastructure EQC 2019-006589-P and the PREVAL 21/07 grant from IDIVAL. We thank Dr. Ilan Davis for his supportive and conceptual contribution to the manuscript. We acknowledge DESY (Hamburg, Germany), a member of the Helmholtz Association HGF, for the provision of experimental facilities. Parts of this research were carried out at PETRA III P62 SAXSMAT beamline. Beamtime was allocated for proposals I-20210691 EC and I-20220193 EC.

## Author Contributions

Conceptualization, A.L.C., S.L, O.M.C., C.S.C., and A.V.V.; methodology, D.M.L., A.N.G., A.P., JMI., HS., G.G., V.M., A.L.C.C., and I.LL.; formal analysis, D.M.L., G.H.O., V.M., A.V.V., and A.L.C.; investigation: protein hybrid synthesis and characterization and analytical studies G.G. and H.S.; investigation: *in vitro* and *in vivo* studies, D.M.L., A.P., H.S., and A.V.V.; investigation: PS-OCT analysis, V.M. and O.M.C.; writing-original draft preparation, D.M.L., G.G., V.M., O.M.C, C.S.C, A.L.C., S.L., C.S.C., and; A.V.V., funding acquisition, A.L.C., C.S.C., and A.V.V. All the authors have read and agreed to the published version of the manuscript.

## Supplementary Figures

**Supplementary Figure 1:**
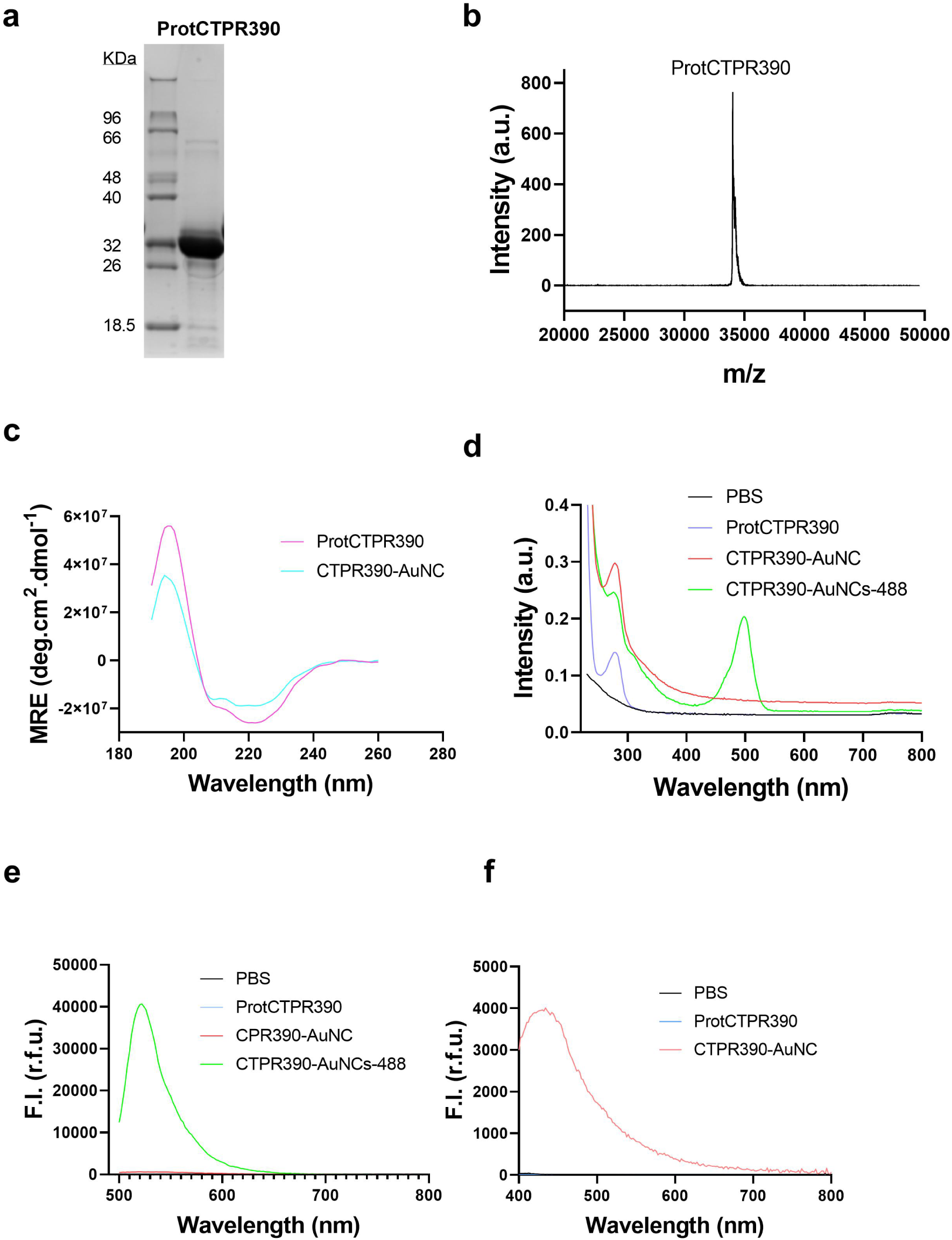
Characterization of CTPR390. **a**) Representative SDS-PAGE showing the purified CTPR390 protein (ProtCTPR390); n = 3 WB belonging to three different batches purified. **b**) MALDI-TOF analysis determined the molecular weight of ProtCTPR390 (34037 Da) through the main peak of the spectrum, reflecting the mass-to-charge ratio (m/z) of detected ions, n = 3 independent assays with 3 replicates. **c**) Circular dichroism (CD) spectra of ProtCTPR390 and the ProtCTPR390 stabilizing Au nanocluster (CTPR390-AuNC) revealed the maintenance of the signature α-helical structure; n = 3 independent assays with 3 replicates. **d**) UV-vis spectrometry highlights variations in light absorption among ProtCTPR390, CTPR390-AuNC (showing typical absorption due to aromatic amino acids at 250-300 nm), and CTPR390-AuNC conjugated to the fluorophore Alexa-488 (CTPR390-AuNC-488) (indicating the presence of Alexa 488 with a peak around 500 nm). CTPR390-AuNC-488 was referred to as CTPR390 in the main text. PBS serves as a control sample; n = 3 independent assays with 3 replicates. **e-f**) Emission spectra showing fluorescence capabilities of CTPR390-AuNC-488 and CTPR390-AuNC compared to ProtCTPR390. The y-axis represents the fluorescence intensity (F.I.) in relative fluorescence units (r. f. u.). **e**) Typical emission spectra of the fluorophore Alexa 488 with excitation wavelength of 460 nm; n = 3 independent assays with 3 replicates. **f**) Specific emissions from CTPR390-AuNC resulting from AuNC excitation at 360 nm. PBS was used as control sample.

**Supplementary Figure 2:**
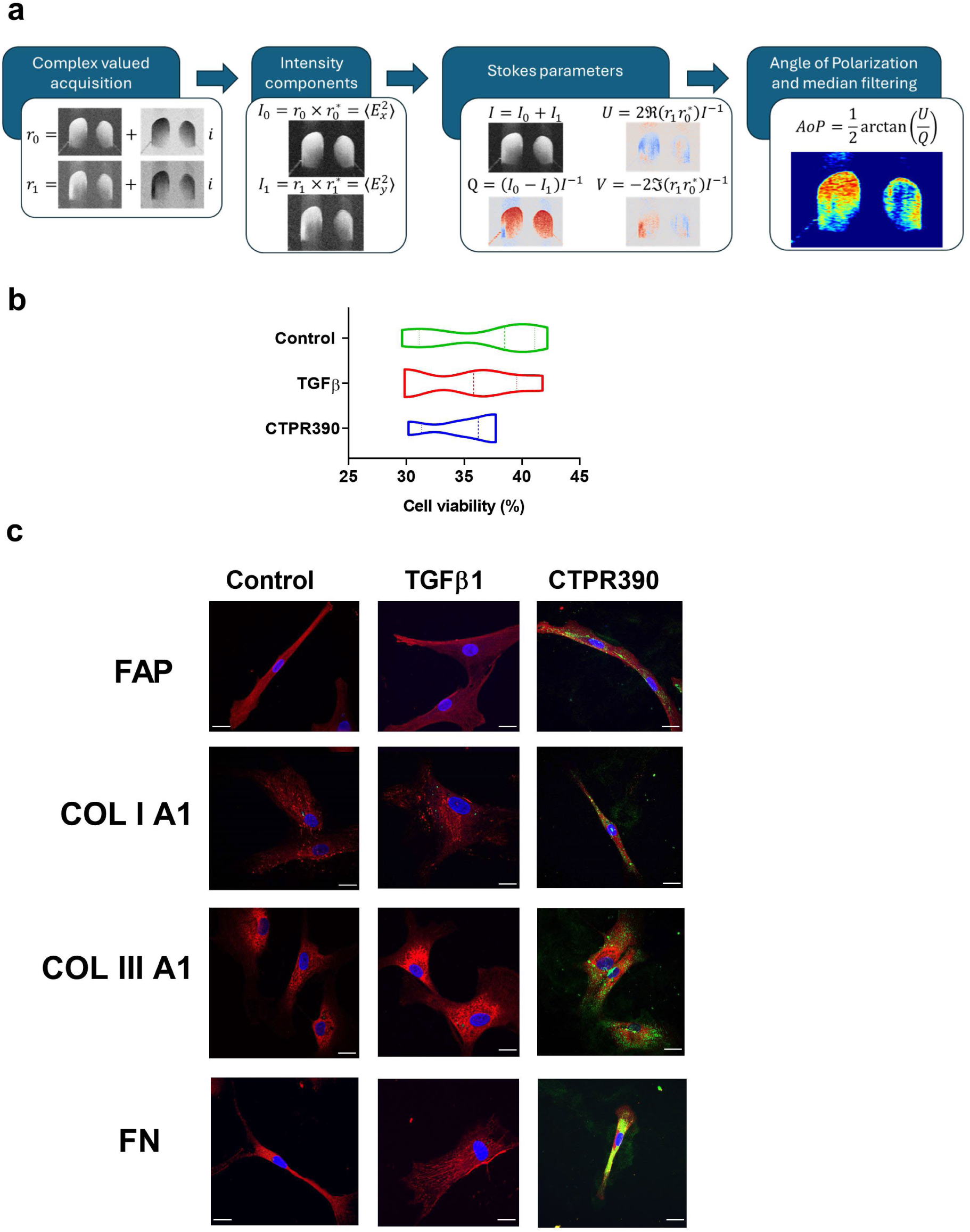
Schematic Illustration of the AoP, calculation, cell viability within ECT, and detection of fibrosis markers in fibroblasts. **a**) Schematic Illustration of the AoP Calculation with the analysis pipeline to obtain the AoP using the PS-OCT system. From left to right, the first blue arrow indicated the changes from the captured values as complex magnitudes to be transformed into intensity components. Second blue arrow marked, that the Stokes parameters are calculated from the captured parallel (I_o_) and perpendicular (I_1_) components of the electric field to calculate the Stokes parameters at all points of the three-dimensional PS-OCT cube (I, U, Q, V). Third blue arrow indicated the step through which the use of Stokes’ U and Q serves to find the AoP. Final data are median filtered with a disk of diameter 3 as a kernel to remove speckle noise. **b**) Violin plot showing the distribution of viable cells within ECT across three experimental groups (Control, TGFβ, CTPR390). Each group includes five samples (n=4-5) and each sample had n = 3 replicates. **c**) Confocal microscopy images of immunofluorescence assays in a 2D culture of human cardiac fibroblasts. Control, TGFβ1 (5 ng / mL), and CTPR390 (1 μM) fibroblasts. The pro-fibrotic markers FAP, COL I A1, COL III A1, and FN were visualized, showing differences in COL III A1 expression. The white scale bars of each panel indicated 10 μm.

**Supplementary Figure 3:**
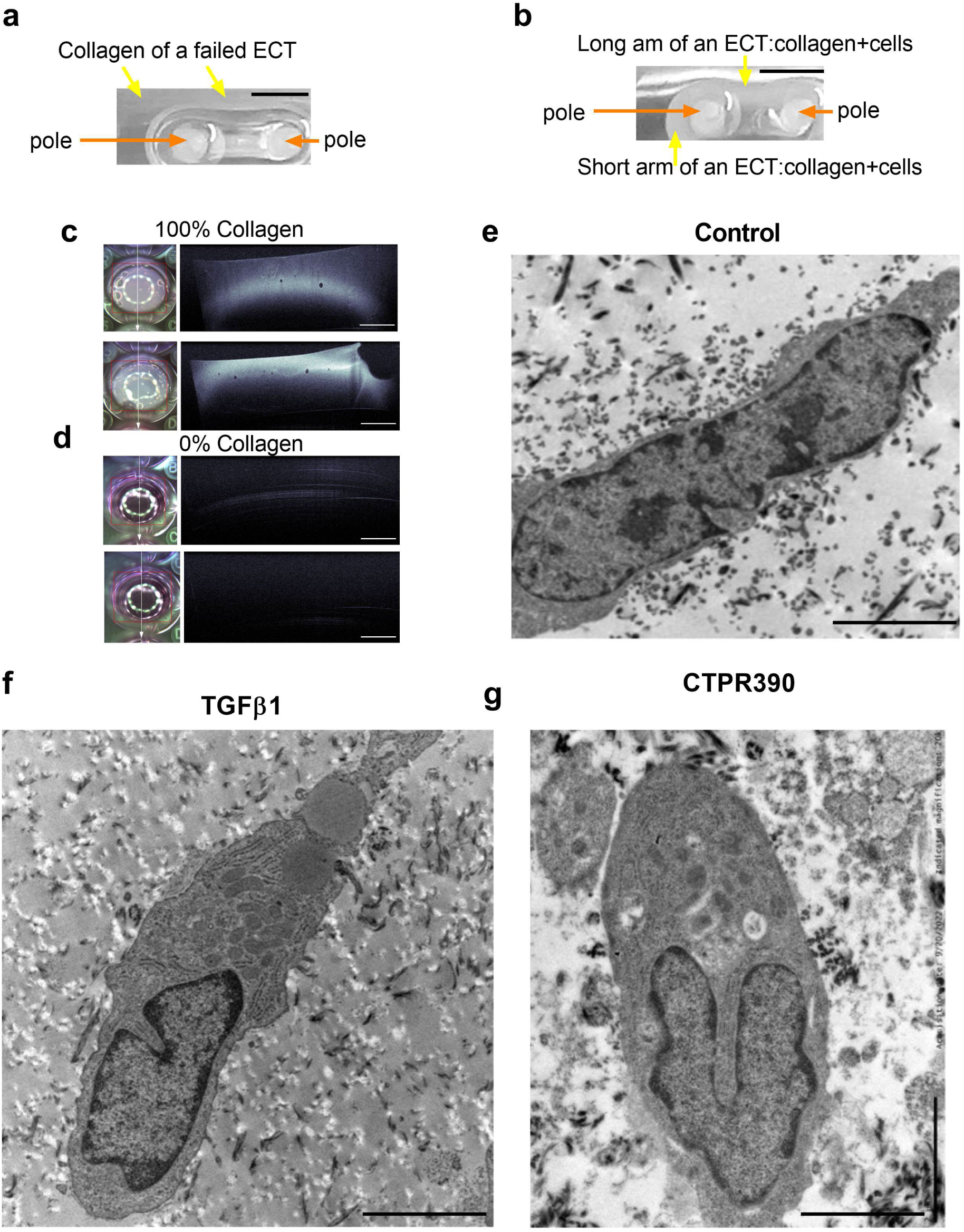
Failed and normal ECT generation and ultrastructure of inner fibroblasts of the ECT. **a-b**) Detailed views of the area surrounding the two poles of an ECT well plate The black scar bars indicated 5 mm. **a**) Close-up view of a failed ECT generated with only collagen and no cells. The viscous material appears as dense white gelatin (yellow arrows) floating away from the two poles (orange arrows). **b**) Close-up view of a matured ECT, showing one short arm and one long arm (yellow arrows) formed with collagen and human cardiac fibroblasts around two poles (orange arrows). **c, d**) Detail of two wells and detection of collagen 100% (**e**), and collagen 0% (**d**), using OCT technique as a dense uniform layer, except for bubbles that appear during solidification. The white scar bars indicated 5 mm.. **e, f, g**) Transmission electron microscopy images of internal fibroblasts showing similar ultrastructural features of Control (**e**), TGFβ1(**f**) and CTPR390 (**g**) ECT on day 13. The black scar bars indicated 2 μm.

**Supplementary Figure 4:**
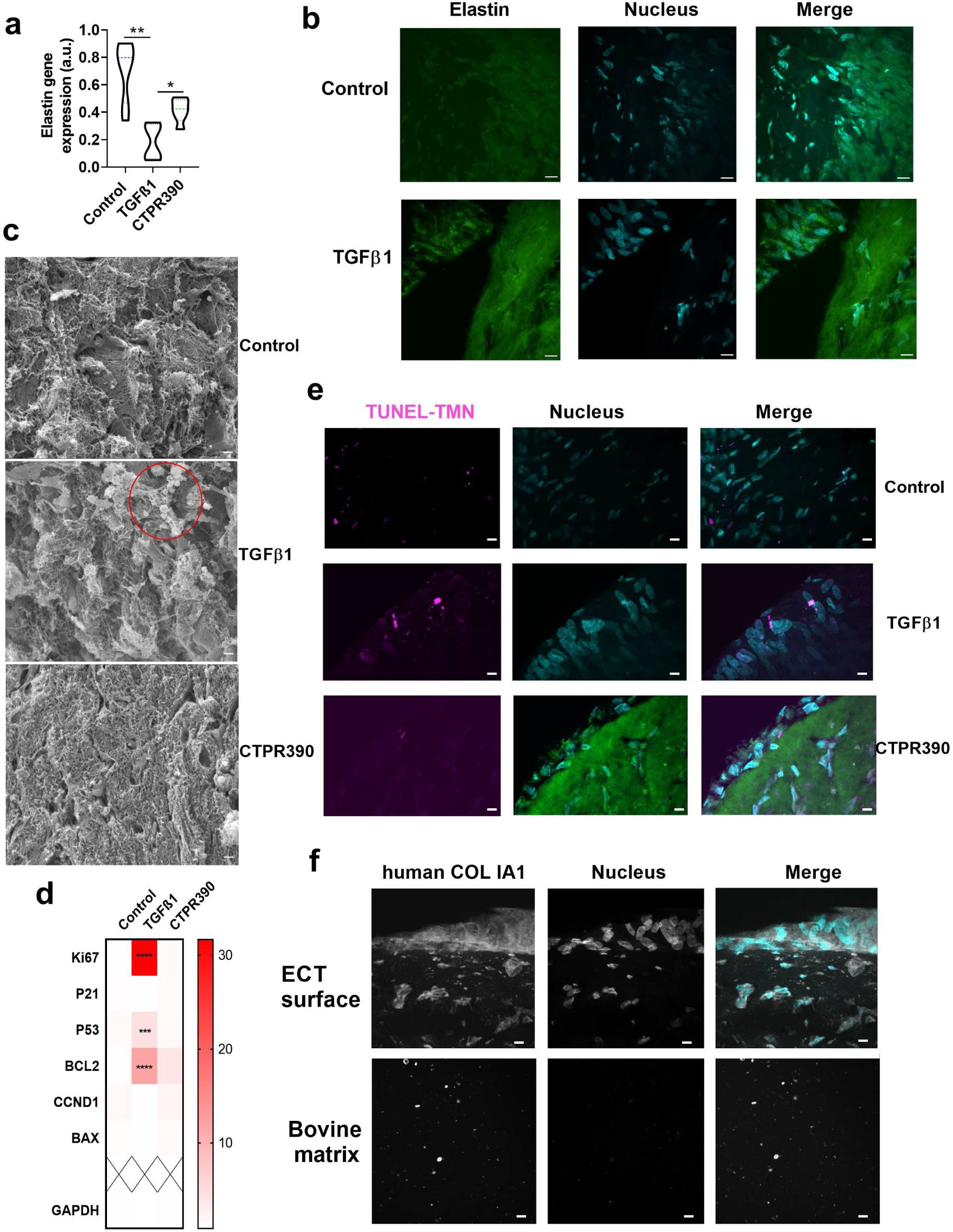
CTPR390 treatment prevents pro-fibrotic ECM activity shown in the TGFβ1 group. Species specificity of the antibody against human COL I A1. **a)** Bar graph of elastin gene expression in Control, TGFβ1 and CTPR390 ECT, n = 5-7 biological samples with 3 replicates. **b)** Confocal images of elastin protein detection in Control, TGFβ1 and CTPR390 ECT; the white scar bars indicated 10 μm. **c)** Representative images of the ECM from the interior of the ECT using scanning microscopy of the long arms of (from top to bottom) Control, TGFβ1 (red open circle indicates apoptotic morphological features) and CTPR390 ECT; the white scar bars indicated 10 μm. **d)** Heatmap illustrating significant differences on genes related to apoptosis (P21, P53, BCL2, CCND1, BAX) and proliferation (Ki67), with darker red representing higher expression (2x over the median) and light reds indicating lower expression (0.5 over the median or lower) in Control, TGFβ1, and CTPR390 ECT groups; n = 6-10 ECT per group with 3 technical replicates of each ECT sample. The p values (*p < 0.05, **p < 0.005, ****p < 0.0001) were determined using a one-way ANOVA with Tukey’s multiple comparison test. **e)** TUNEL assay marking more apoptotic areas of the ECM in TGFβ1 compared to Control and CTPR390 groups; The white scale bars indicated 10 μm. **f)** Confocal images showing human COL IA1 specificity on human collagen vs bovine collagen. Detection of human collagen (COL IA1 in white) generated by human fibroblasts (nuclei DAPI staining in cyan) and merge of both images of the ECT (upper images) and absence of detection in bovine matrix without cells (lower images) used as basal collagen matrix of the ECT. The white scale bars indicated 10 μm.

**Supplementary Figure 5:**
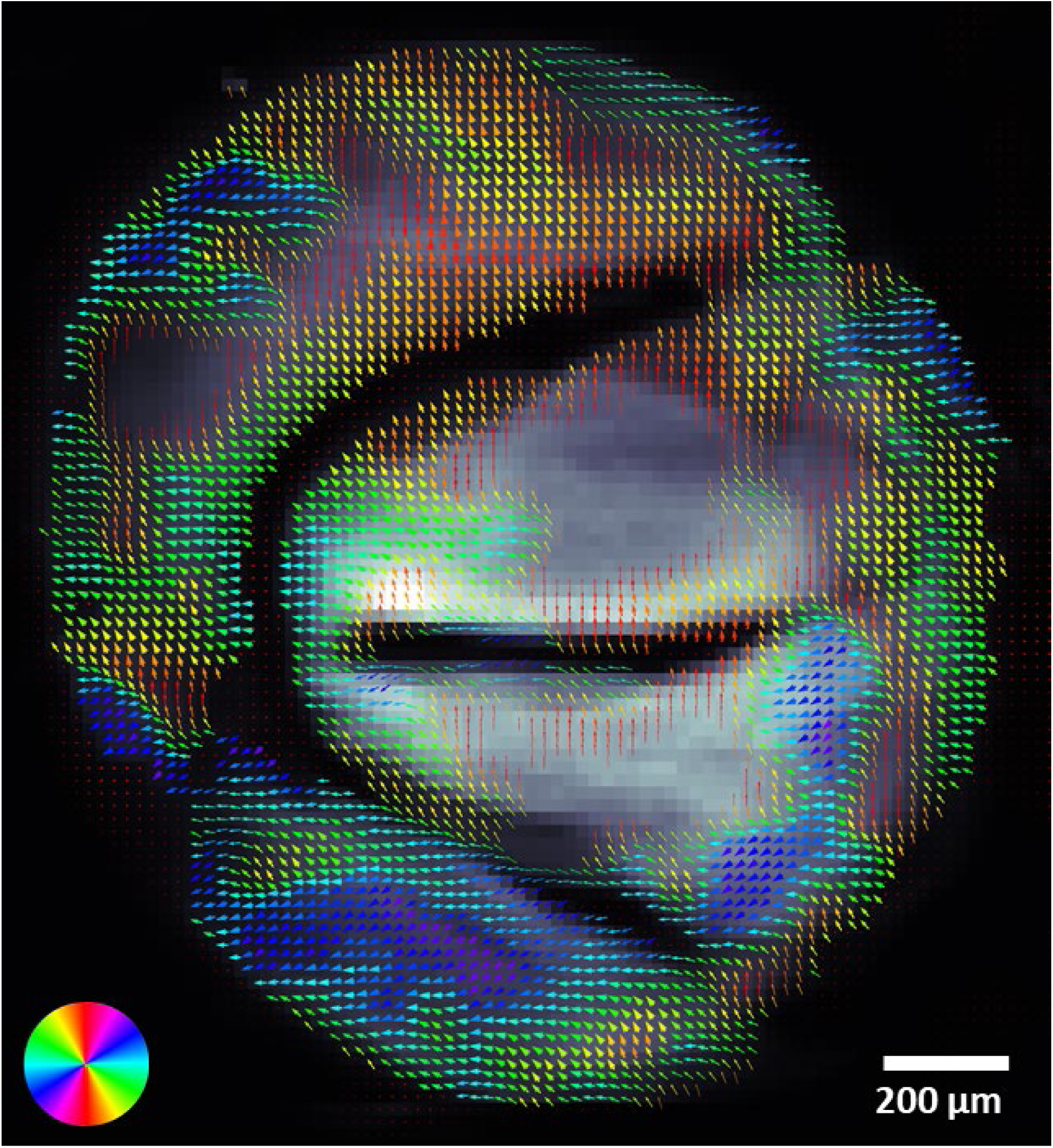
Organization of collagen produced by human fibroblasts on the surface of Control ECT. Map of the preferential orientation and degree of orientation of collagen in control ECT. White bar indicates 200 μm. Preferential of orientation in each 20 x 20 µm^2^ pixel is indicated by the direction of the arrow in each of the pixels of the figure and the colour coded disc. Degree of orientation in each 20 x 20 µm^2^ pixel is indicated by the thickness of the arrows in each of the pixels of the figure. The white scale bar indicated 200 μm.

**Supplementary Figure 6:**
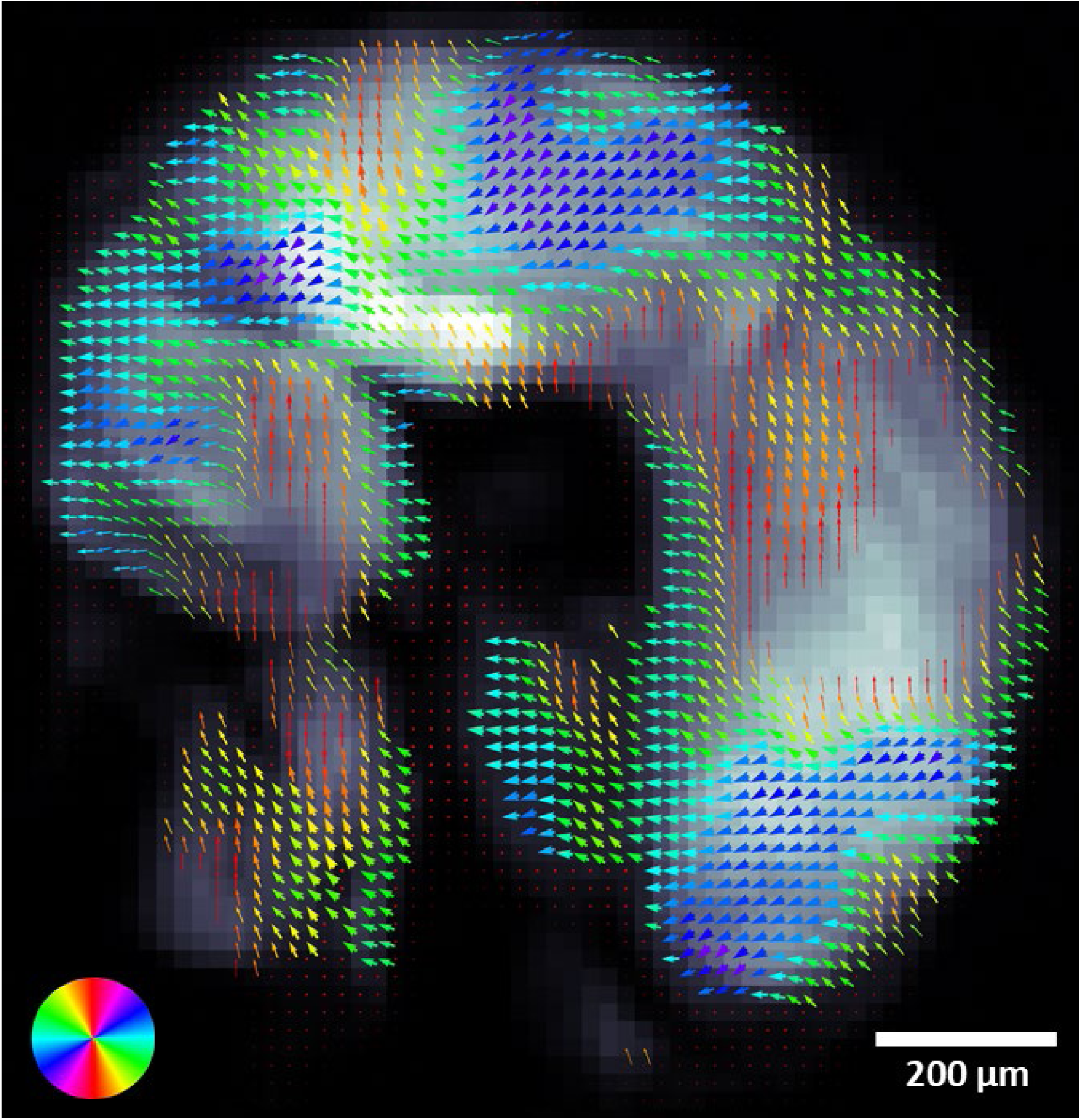
Organization of collagen produced by human fibroblasts on the surface of TGFβ1 ECT. Map of the preferential orientation and degree of orientation of collagen in TGFβ1 ECT. White bar indicates 200 μm. Preferential of orientation in each 20 x 20 µm^2^ pixel is indicated by the direction of the arrow in each of the pixels of the figure and the color coded disc. Degree of orientation in each 20 x 20 µm^2^ pixel is indicated by the thickness of the arrows in each of the pixels of the figure. The white scale bar indicated 200 μm.

**Supplementary Figure 7:**
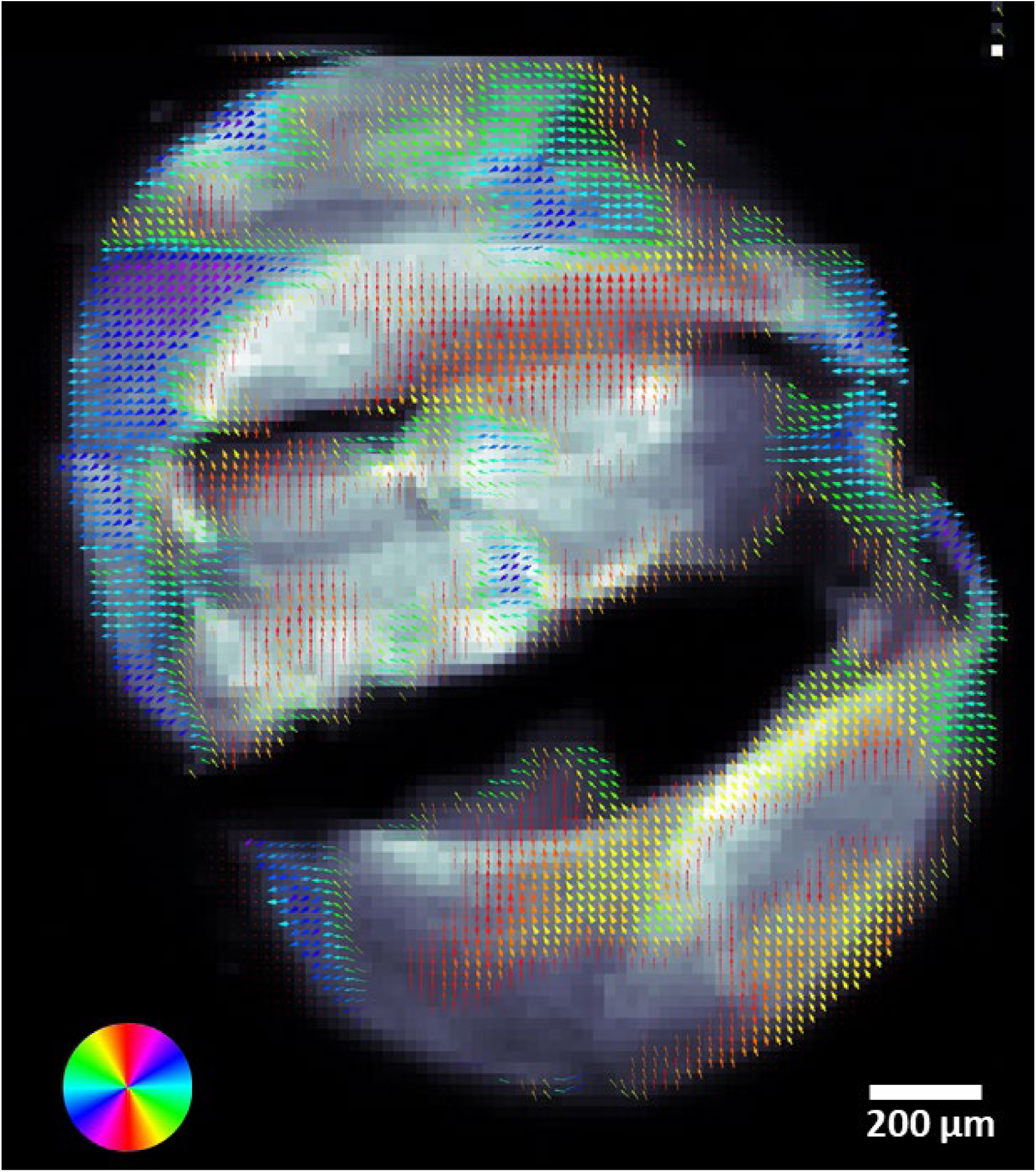
Organization of collagen produced by human fibroblasts on the surface of CTPR390 ECT. Map of the preferential orientation and degree of orientation of collagen in CTPR390 ECT. White bar indicates 200 μm. Preferential of orientation in each 20 x 20 µm^2^ pixel is indicated by the direction of the arrow in each of the pixels of the figure and the color coded disc. Degree of orientation in each 20 x 20 µm^2^ pixel is indicated by the thickness of the arrows in each of the pixels of the figure. The white scale bar indicated 200 μm.

